# Species-associated bacterial diversity increases along a gradient of habitat degradation

**DOI:** 10.1101/2024.01.04.574207

**Authors:** Elina Hanhimäki, Susanna Linna, Camila Beraldo, Mikael Englund, Uxue Rezola, Pedro Cardoso, Rose Thorogood, Marjo Saastamoinen, Anne Duplouy

## Abstract

Alterations of microbial communities have evident impacts on development, digestion, fecundity, metabolism, immunity, and diverse other biological functions of their hosts. Yet, the factors affecting microbial communities associated with wild species often remain uncharacterized. For example, the impact of the host’s habitat degradation due to anthropogenic activities has received little attention, which contrasts with the large literature showing how such habitat degradation is at least partly responsible for the on-going global patterns of macro-biodiversity erosion. Here, we use metacommunities of herbivorous insect species specialized in feeding on *Plantago lanceolata* in the fragmented landscape of the Åland Islands, Finland, as a model system to test whether and how bacterial communities associated with wild species change along a gradient of habitat degradation. We evaluated microbial species diversity and community composition in two sympatric insect species sampled from local meadow habitats with various levels of human disturbance within or around these habitats (e.g. forests, roads, agriculture fields, buildings). Counter to our expectations, we found that bacterial diversity can increase with habitat degradation, with individuals from more degraded habitats hosting more rare bacterial species. In contrast, as the dominant microbial species remain similar across habitats, the community composition and function of the microbiota persist under habitat degradation. In this system, the strength of human activities might induce changes in habitat heterogeneity rather than changes in overall habitat quality, thus allowing local insects to encounter and host more rare microbes rather than trigger local microbial extinction.

## Introduction

Deterministic processes drive assembly patterns of resident microbial communities, while stochastic processes are more likely to result in the acquisition or loss of random and transient microbial species (Bénard et al 2020, Moran 2007, Unzueta-Martínez et al 2021). In either case, alterations in the functional diversity of microbiota can impact the development, digestion, fecundity, metabolism, immunity, and/or diverse other biological functions in hosts (Engel and Moran 2013, Jing et al 2020), but see (Duplouy et al 2020, Hammer et al 2017). Therefore, changes in microbiota may scale up to influence the resilience of macrobiota and the wide range of community interactions and ecosystem services that they provide (Thorogood et al, 2023). Nevertheless, while a large literature illustrates that habitat degradation is at least partly responsible for on-going global patterns of erosion in macro-biodiversity (Borges et al 2020, Decaëns et al 2018, Horváth et al 2019), the impacts of habitat degradation experienced by the host on their associated microbial biodiversity have received little attention (Hom and Penn 2021).

Habitat degradation is a multifaceted umbrella term, which is generally associated with negative impacts on biodiversity through decline of the local species richness or changes in community composition leading to biotic homogenization (Borgella et al 2001, García-Martínez et al 2017, Kumar and O’Donnell 2007),(and see reviews by Cardoso et al 2020, Soh et al 2019). For example, in their global analysis of the effect of land-use change on the megadiverse group of the rove beetles, Méndez-Rojas et al. (2021) showed that species density and richness decrease with increasing transformation of the beetles’ natural habitats into agricultural fields, while Uhl et al. (2021) demonstrated that decrease in habitat quantity across the landscape negatively affect the functional diversity of moths in Italian forest reserves. In contrast, fragmentation per se (i.e. breaking of the habitat into smaller fragments without total surface loss) has been argued to possibly promote local biodiversity due to the overall higher levels of habitat heterogeneity (Fahrig 2003, Riva and Fahrig 2023). Although the debates on how qualitative and quantitative features of the habitat may affect macro-biodiversity are the theme of a large literature, little is known on how these features might affect the micro-communities associated with macro-species. On one hand, under environmental change we might anticipate that variation in the microbiota associated with species should reflect active selection or purging of microbes beneficial or detrimental to the host’s abilities to tolerate or resist stresses (Fagundes et al 2012, Houwenhuyse et al 2021, Lemoine et al 2020, Tougeron and Iltis 2022, Wernegreen 2012). However, it is also plausible that microbial changes could simply reflect changes in the environmental microbiota, which the host encounter and collect at random through dispersing and feeding (Hannula et al 2019, Mason et al 2021).

A bedrock poor in nutrients but locally covered in rich sediments supports the different habitat types observed across the Åland Islands, a Finnish archipelago in the Baltic Sea (Haila et al 1980). The islands offer areas covered by extremely patchy managed forests, meadows or peatlands, while they also experience growing agricultural and residential development pressures (EuropeanCommission 2022, Haila et al 1980). In this particular environment, a scattered network of over 4,400 meadows and pastures, colonized by the common ribwort plantain, *Plantago lanceolata*, have been geolocated at the interface of agricultural and natural landscapes, and systematically surveyed since 1993 (Nieminen and Vikberg 2015, Ojanen et al 2013). These meadows are habitat patches for many species, including the Glanville fritillary butterfly *Melitaea cinxia* (Linnaeus, 1758) and the weevil *Mecinus pascuorum* (Gyllenhal, 1813), two specialized herbivores of *P. lanceolata* (Nieminen and Vikberg 2015, Ojanen et al 2013). Each meadow differs from the next in many different quality aspects, including their surface area (i.e. the median meadow size is of 0.06 hectares, Hanski et al 2017), as well as the level of habitat degradation due to different human activities within the meadows themselves, and in the landscape surrounding each meadow (Schulz et al 2019).

Recent studies have described the gut microbiota of *M. cinxia* caterpillars as transient (Duplouy et al 2020, Minard et al 2019), and possibly only representative of their environment. These microbial studies are consistent with studies in other Lepidoptera species (Hammer et al 2017), and supplement the wealth of ecological, genetic, and genomic data already available for this butterfly species (Kahilainen et al 2018, Smolander et al 2022, van Bergen et al 2020). In contrast, the details of the relationship between *M. pascuorum* weevils and their associated microbiota remains unknown. Nonetheless, based on the literature, these weevils are likely to carry a resident microbiota, dominated by few bacterial symbionts with important mutualistic functions for their host fitness (Anbutsu et al 2017, Hsiao and Hsiao 1985, White et al 2015). This system thus offers an opportunity to test whether habitat degradation, from meadows to agricultural and residential habitats, have impact on the microbial communities associated with these two sympatric insect species.

Here, we used metabarcoding techniques to independently characterize the bacterial microbiota associated with *M. cinxia* butterfly and with *M. pascuorum* weevil specimens, which were collected from meadows scattered across the Åland Islands. We expected that the quality of the host’s habitat would positively corelate with bacterial species richness in these insects, and that bacterial community composition would differ between insects from small and large meadows, and/or with the level of Anthropogenic disturbances within the meadows, or within the landscape. In these conditions, insect- associated microbial communities could not only act as indicators of their hosts’ health, but also of the quality of the hosts’ habitat.

## Materials and methods

### Samples

In total, we used a set of 99 *M. cinxia* caterpillars previously collected from 38 meadows from seven communes (Eckerö, Finström, Geta, Hammarland, Jomala, Lemland and Sund), and a set of 125 *M. pascuorum* weevils collected from 16 meadows from eight communes (Finström, Geta, Hammarland, Jomala, Lemland, Lumparland, Sund and Vårdö) (Fig. 1) within the Åland Islands. The *M. cinxia* caterpillars were collected in September 2012 as they diapause gregariously within overwintering silk nests (Ojanen et al 2013). About one third of the wild collected *M. cinxia* caterpillars are naturally parasitized by the specialized parasitoid wasp, *Hyposoter horticola* (Gravenhorst, 1829) (Duplouy et al 2015, van Nouhuys et al 2012, van Nouhuys et al 2016) but detection of the parasitoid wasp at this stage of development is only possible through PCR or dissection (see below). The weevil *M. pascuorum* were collected in June 2020, while conspicuously feeding on the host plant or laying eggs on the host plant seeds (Nieminen and van Nouhuys 2017). Adult weevil species are not known for being attacked by parasitoid wasps. All *M. cinxia* caterpillar and *M. pascuorum* weevil specimens were collected alive, individually placed in a tube labelled with a unique barcode and stored at –20°C until further manipulated. The origin and metadata associated with each specimen, including characteristics of the meadows, can be found in Table S1.

**Figure 1.**
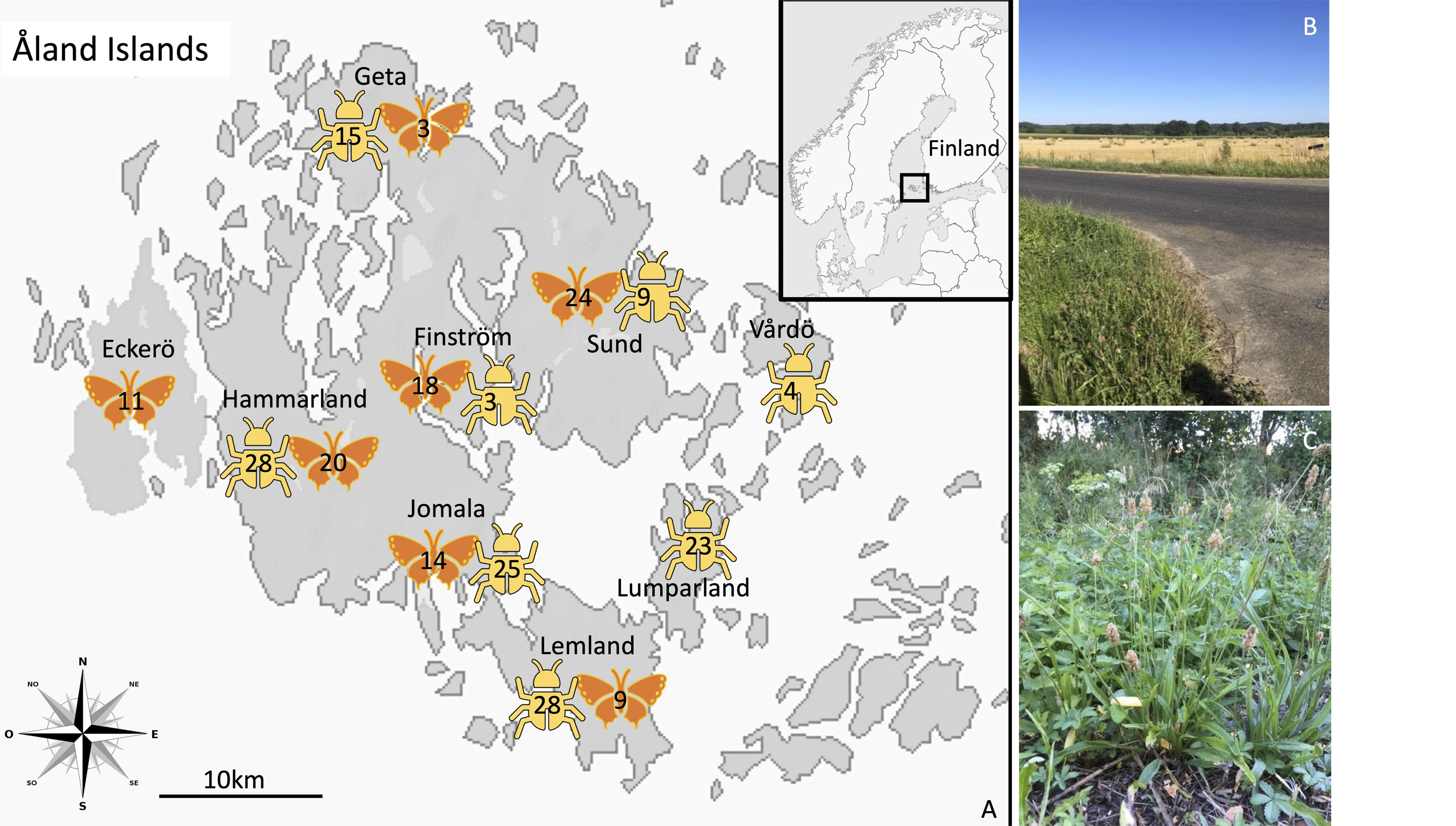
Sampling locations for *M. cinxia* caterpillars (orange symbols) and *M. pascuorum* weevils (yellow symbols) across the Åland Islands, Finland. Numbers on top of the symbols indicate the number of specimens collected within each sampling commune and species. (Right top): a meadow within degraded habitat type, (Bottom right): host plant *Plantago lanceolata* with flowers.

To test the effect of habitat degradation on the microbiota of the two host species, we identified the types of land-use within and around a 10-meter buffer area of each sampled meadow. The different land-use types were previously described by Schulz et al (2019) and represent 13 land-use categories, either associated with natural habitats (e.g. beaches, rocks, meadows, forests, streams), or associated with habitat degradation due to anthropogenic activities (e.g. agricultural fields, roads, built areas). Each meadow was assigned two values of habitat degradation, as the proportion of degraded versus natural habitats (1) within the meadow, and (2) within the 10-meter buffer surrounding the meadow. Land-use within the meadow captures direct aspects of the meadow quality, while the land-use within the 10-meter buffer surrounding each meadow capture wider overall landscape degradation (Schulz et al 2019), which can affect the microclimatic conditions or colonization of the meadow. Similarly, as pressures from land-use changes due to anthropogenic activities within and around each meadow are likely to increase as the size of the meadow decreases, we collected the surface area for each meadow (m^2^) from Ojanen et al (2013) and Hanski et al (2017) (Table S1).

### Molecular work: DNA extraction and PCRs

All specimens were individually washed by plunging them in a 1X PBS bath for a few seconds, and dissected under a sterile laminar flow cabinet to avoid environmental contamination at early stage of the sample preparation. We dissected out the content of the gut of each *M. cinxia* caterpillar, while we used entire adult *M. pascuorum* weevils for individual DNA extractions using a Qiagen DNeasy Blood and Tissue kit (Qiagen, Germany) following manufacturer’s protocol. Seven blank samples (sterile water samples; two per 96 well PCR plate) were similarly processed as controls for contamination across the extraction protocol. Upon dissection, we noticed that 18 of the 99 *M. cinxia* caterpillars looked different (rigid bodies and little internal material), which may suggest these specimens were already deceased before preservation. However, as this did not affect any of the following analyses this observation was not further investigated. We tested for parasitism by the parasitoid wasp *H. horticola* within the *M. cinxia* caterpillars by amplifying a section of the protein-b11 from the wasp-integrated ichnovirus, using the primer pair 34F-463R (Duplouy et al 2015).

The V5-V6 hypervariable region of the conserved bacterial 16S rRNA genes was amplified by PCR, using the modified 784F/1061R primers (Andersson et al 2008). The amplification was performed on BioRad thermal cycler with a program of 5 min at 95°C, followed by a total of 40 cycles at 95°C for 40 s, 54.2°C for 45s, 72°C for 40s and ending with the final extension at 72°C for 7 minutes. All specimens were amplified in duplicate using 3 µl of DNA extract (Duplouy et al 2020, Minard et al 2019). The duplicates were pooled and sequenced using a MiSeq v.3. illumina Sequencing platform at the Finnish Institute for Molecular Medicine, FIMM.

### Metabarcoding data processing

Sequence data was demultiplexed at the FIMM sequencing centre. Paired-end reads were processed with QIIME2 v.2021.2 (Bolyen et al 2019) separately for *M. cinxia* and *M. pascuorum* datasets but using the same pipeline. Primer sequences were removed using the QIIME2 plugin cutadapt v.2021.2.0. Read-pairs were joined, quality trimmed and denoised using DADA2 v.2021.2.0 (for *M. cinxia*, forward read trimmed at 224 and reverse read trimmed at 135; for *M. pascuorum* forward read trimmed at 245 and reverse read trimmed at 195). Taxonomy was assigned to Silva 138 database (Quast et al 2013) using the q2-feature-classifier plugin (Bokulich et al 2018). Amplicon sequence variants (ASVs) that were not assigned to any bacterial phyla, or that were classified as mitochondria or chloroplasts, were removed. Next, we used the R v.4.0.4 package decontam v.1.10.0 (with the prevalence-method using a threshold of 0.5)(Davis et al 2018) to remove any procedure contaminant from the data based on the protocol controls (as described in Duplouy et al 2020, Minard et al 2019), after which a midpoint rooted tree was produced.

### Statistical analyses

All analyses were done separately for the two species. All statistical tests were completed using QIIME2 v.2021.2 (Bolyen et al 2019) and R v.4.0.4 (RCoreTeam 2020) using the packages ggplot2 v.3.3.2 (Wickham 2016), vegan v.2.5.7 (Oksanen et al 2020), and indicspecies v.1.7.12 (De Cáceres and Legendre 2009). We tested our variables for normality using Shapiro-Wilk normality test (Shapiro and Wilk 1965). We constructed principal coordinate analysis (PCoA) plots using phyloseq v.1.34.0 0 (McMurdie and Holmes 2013) to visualize any patterns of clustering throughout the habitat disturbance gradient.

#### a) Species diversity (alpha-diversity) and Indicator species

To test the effect of the habitat degradation, within the 10-meter buffer area surrounding each meadow or within the meadow itself, on the alpha-diversity of the microbiota of *M. pascuorum* or *M. cinxia*, we used linear models (with Shannon index) and negative binomial regression models (for observed ASVs count) including the level of habitat degradation (within the meadow or within a 10-meter buffer) and the village from which the sample was collected as predictors to the models. Additionally, we added meadow size and for *M. cinxia* only, parasitism by *H. horticola,* as additional explanatory variables to the models. The meadow size and parasitism status were however removed from the models to avoid overfitting, after the AIC model selection test showed adding them increased the complexity of the model without a significant improvement of its performance.

For each host species, we used the indicspecies v.1.7.12 package (De Cáceres and Legendre 2009) with the IndVal index and 9999 permutations to identify indicator ASVs for habitat degradation. Indicator species are ASVs that are either less or more abundant in either sample category compared. As the indicspecies model does not accept continuous variables, we categorized the meadows into two groups: (1) meadows surrounded by a 10-meter area with no to little degradation (proportion of human activities < 0.25); and (2) meadows surrounded by a 10-meter buffer area, of which more than a quarter is affected by human activities (proportion of human activities ≥ 0.25).

#### b) Species community composition (beta-diversity) and predictability

To assess changes in the composition of microbiota of *M. cinxia* and *M. pascuorum*, we visualized the data using a principal coordinate analysis (PCoA) and calculated permutational multivariate analyses of variance (adonis PERMANOVA with 999 permutations) for Bray Curtis dissimilarity (Bray and Curtis 1957) and Unweighted UniFrac (Lozupone et al 2011) including the level of habitat degradation (within the meadow or within a 10-meter buffer), the meadow size, and the sampling village as explanatory variables using adonis2 function in vegan v.2.5.7 (Oksanen et al 2020). To visualize the five most important bacteria affecting sample clustering ordination biplots were assessed for both Bray Curtis and unweighted UniFrac. Parasitism status of the *M. cinxia* caterpillars, which showed no statistical effect on the model, was removed from the model to avoid overfitting.

Furthermore, we examined how predictable, and thus distinguishable, the taxonomic profiles of the microbial communities associated with either *M. cinxia* or *M. pascuorum* were throughout the gradient of habitat degradation, by using a random forest regression model with 100 trees in QIIME2 with plugin q2- sample-classifier (Bokulich et al 2018).

### Predicted functionality

Finally, to examine changes in the predicted functional composition of the microbiota of *M. cinxia* or *M. pascuorum* along the gradient of habitat degradation, we used the QIIME2 v.21.11 plugin q2-PICRUSt2 (Douglas et al 2020). This plugin uses the pathway abundance method and the EPA-ng placement tool (Barbera et al 2019) to produce and plot a Bray Curtis matrix with predicted functional pathways. We examined any patterns of specimen clustering in R, using adonis2 model (PERMANOVA with 999 permutations) including the level of habitat degradation within each meadow or in the 10-meter buffer surrounding each meadow, the meadow size, and the sampling village as explanatory variables.

## Results

For the butterfly microbial dataset, a total of 2,553,069 sequences remained after sequence cleaning and decontamination (8,741,919 original read pairs), which provided for 6,836 amplicon sequence variants (ASVs) with an average of 25,789 sequences and 373 ASVs per sample. This dataset was normalised with rarefaction to 10,293 sequences, without leaving out any samples. The most abundant phyla associated with the gut of *M. cinxia* caterpillars were Proteobacteria, Bacteroidota and Actinobacteria (Fig. 2A), whereas the most abundant genera were *Yersinia*, *Sphingomonas* and *Hymenobacter* (Fig. S1A).

**Figure 2:**
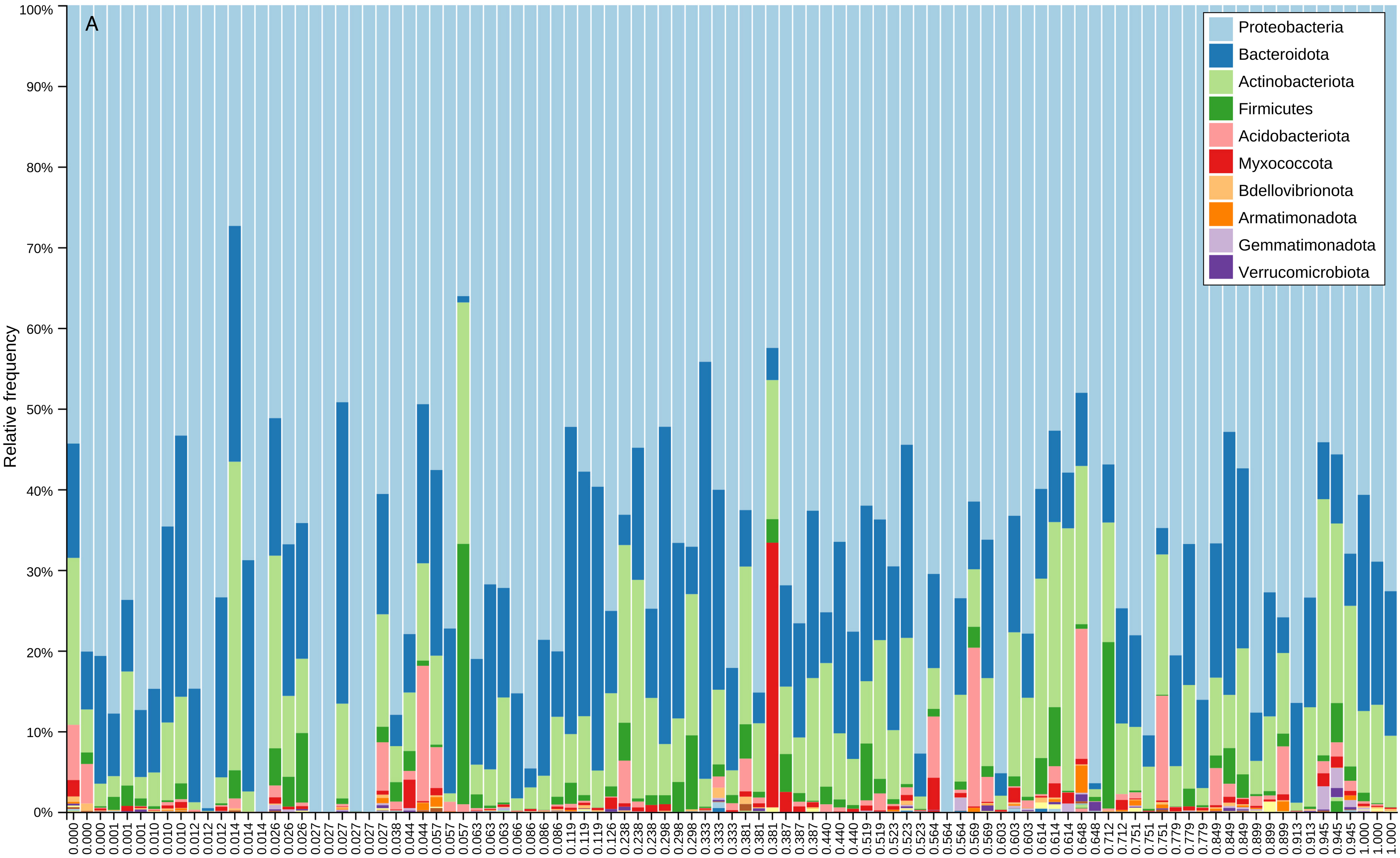
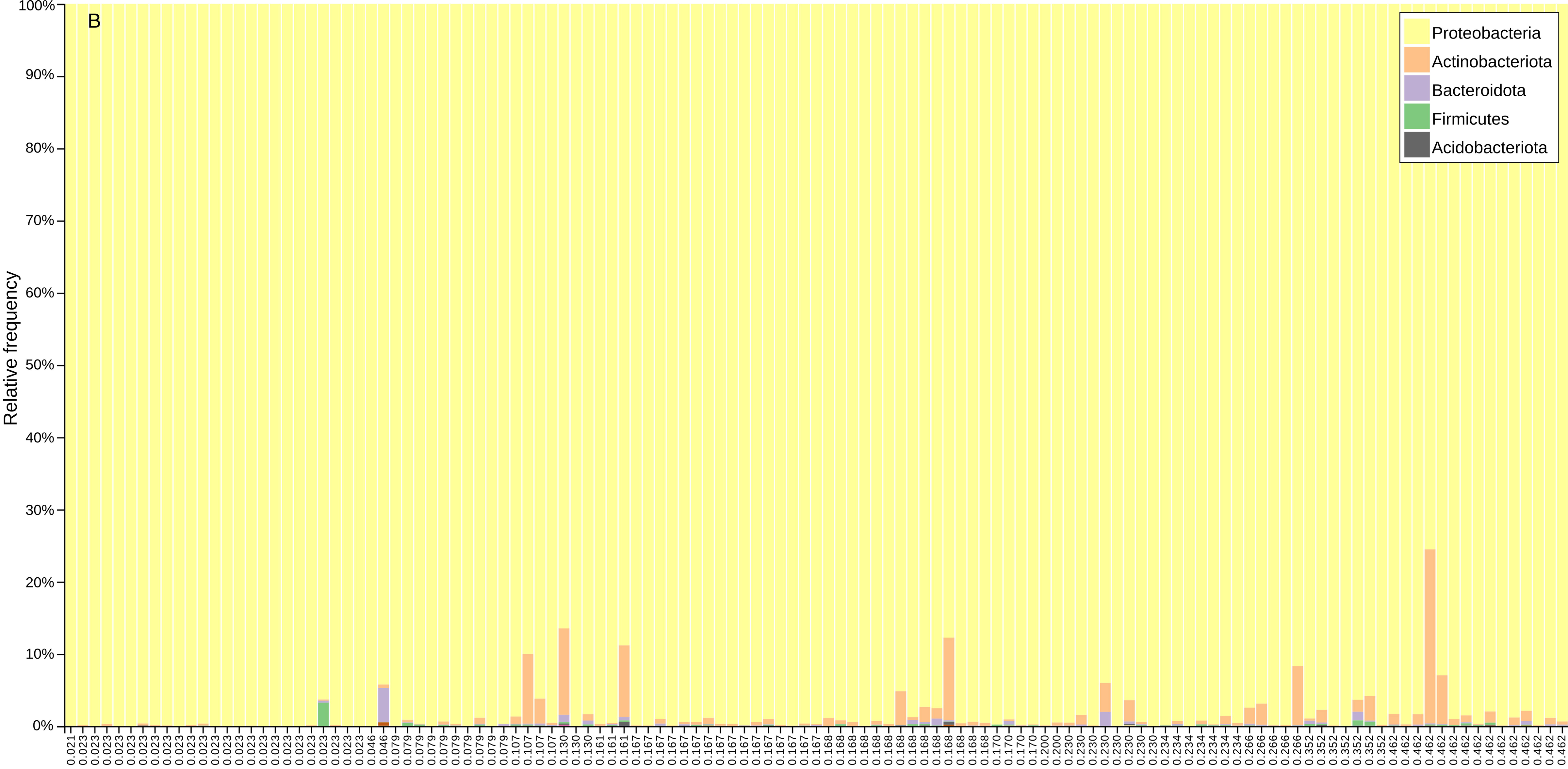
Relative abundance of bacterial phyla making the microbiota colonizing (A) the gut of *M. cinxia* caterpillars, and (B) the full body of *M. pascuorum* weevils.

For the weevil microbial dataset, a total of 3,018,930 sequences remained after sequence cleaning and decontamination (14,849,912 original read pairs), which provided for 3,105 ASVs, with an average of 54,881 sequences and 2,227 ASVs per sample. This dataset was rarefied to 28,936 sequences with no sample loss. The most abundant phyla associated with *M. pascuorum* were also Proteobacteria, Actinobacteria and Bacteroidota (Fig. 2B), whereas the most abundant genera were three bacterial symbionts: *Rickettsia*, *Sodalis* and *Wolbachia* (Fig. S2).

The random forest regression model could not predict the microbiota of the butterfly *M. cinxia* under habitat degradation in the 10m buffer surrounding each meadow (R2 = 0.001; mean squared error = 0.008; *p*-value = 0.908; Fig. 3A, Table S2); but a similar model accurately predicted the bacterial profile of the weevil *M. pascuorum* (R2 = 0.748; mean squared error = 0.11; *p*-value=2.47E-08; Fig. 3B, Table S2).

**Figure 3.**
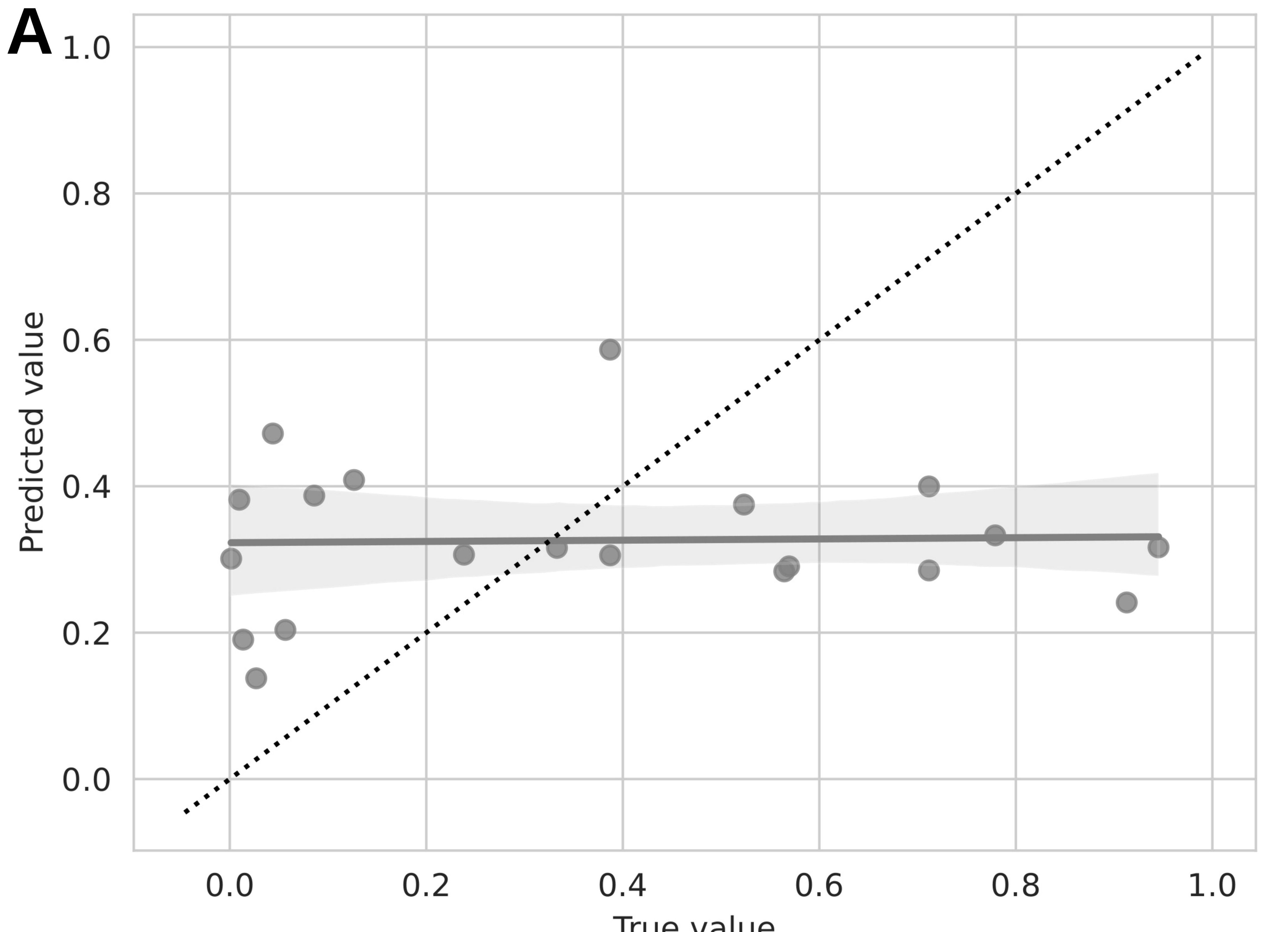
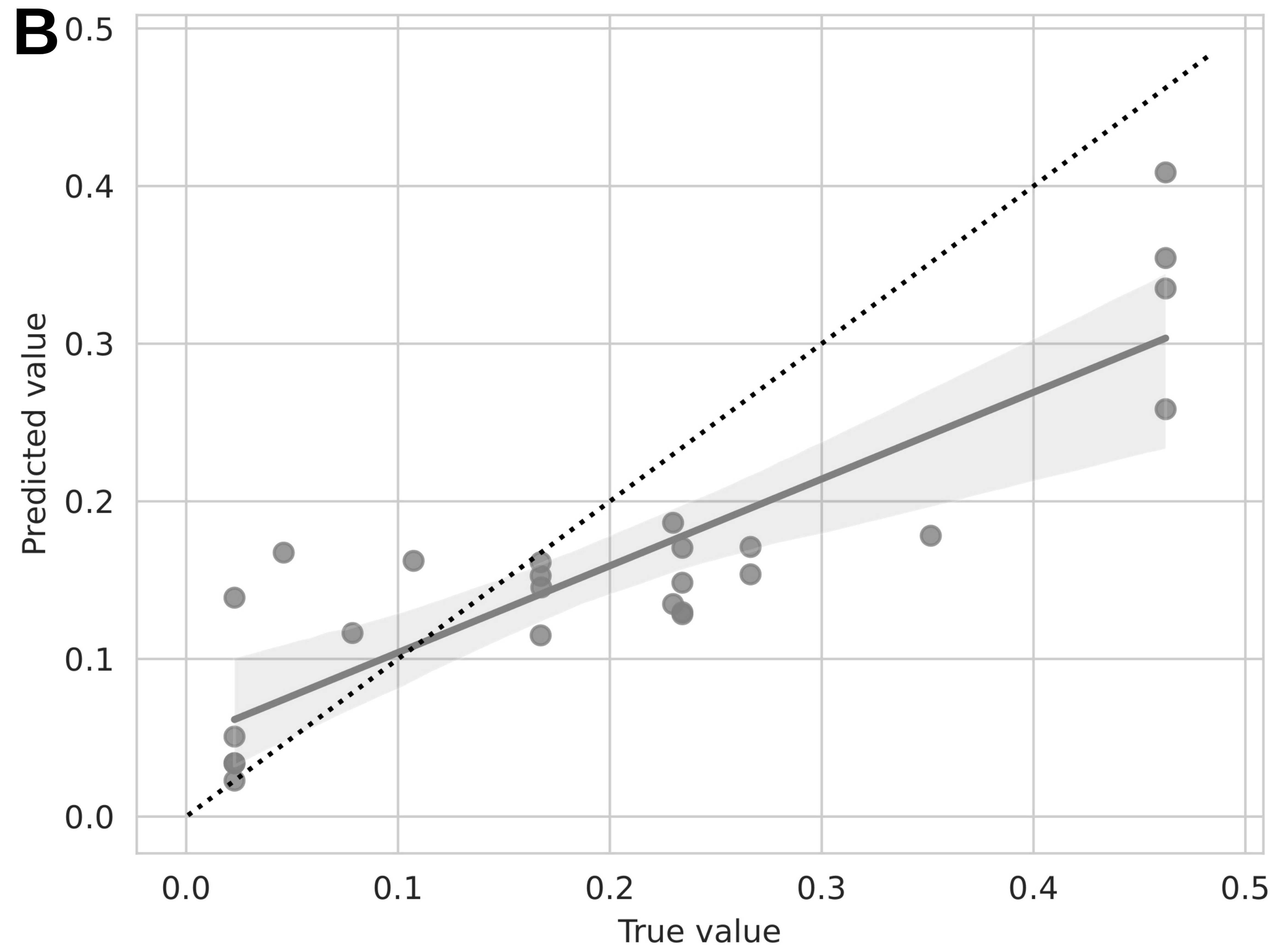
Scatterplots with linear regression lines with 95% confidence intervals presenting the accuracy results of the random forest q2-classifier for microbiota samples from (A) *M. cinxia* or (B) *M. pascuorum*. The dotted line represents the ideal 1:1 ratio between predicted and accurate values of microbiota composition.

### Effects of habitat degradation on bacterial diversity (alpha-diversity) and Indicator species

For the butterfly *M. cinxia*, the level of degradation within habitat or of degradation in the 10m buffer surrounding each meadow had no effect on the observed ASV count, nor on the Hill-Shannon diversity of the microbiota (*p*-values > 0.05; Fig. S3A&B; Fig. 4A&B, Table S3). The focal meadow size and parasitism by parasitoid had no effect on the observed ASV count, nor on the Hill-Shannon diversity of the microbiota (*p*- values > 0.05), and were thus removed from the model as explained in the method section.

**Figure 4.**
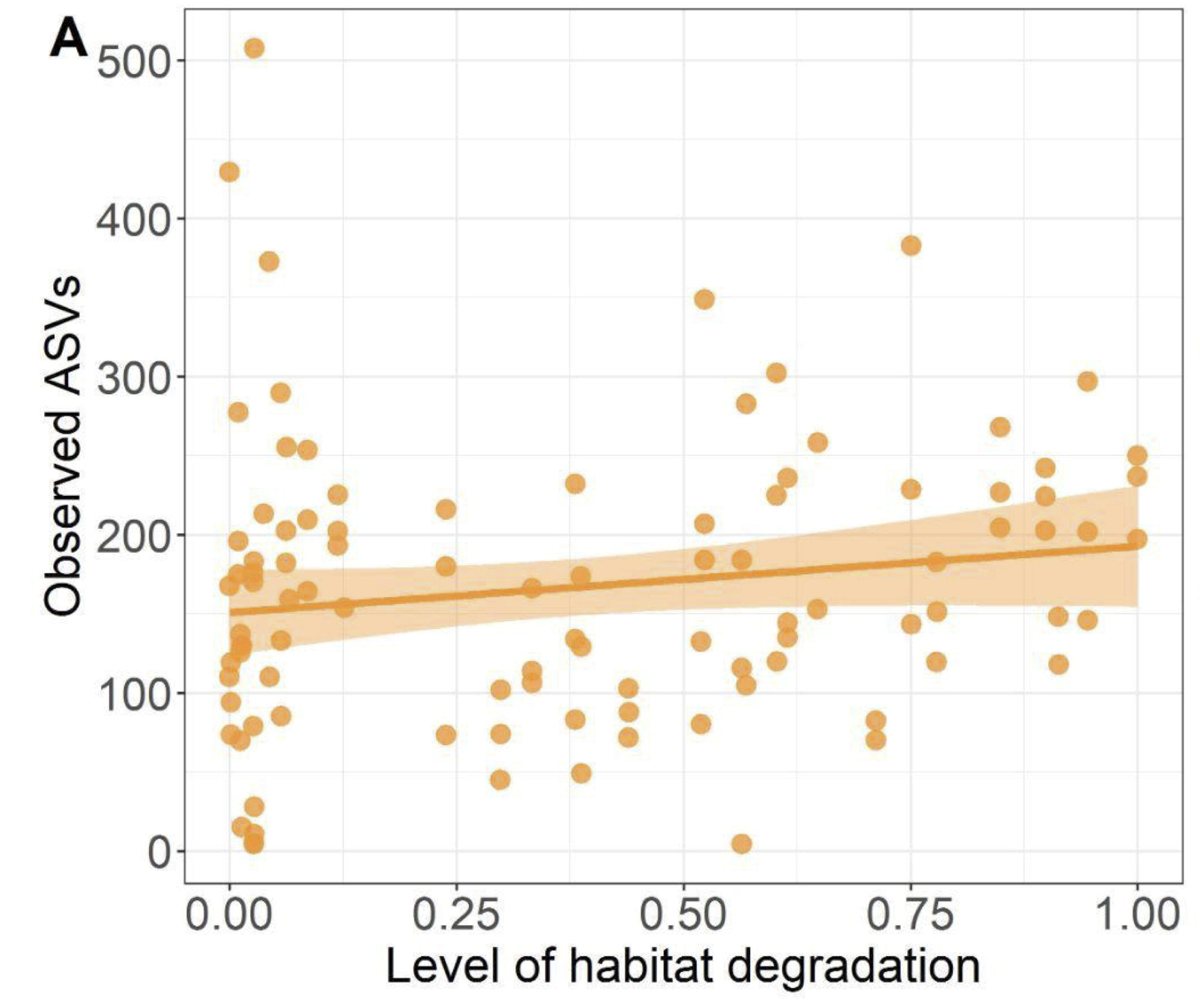
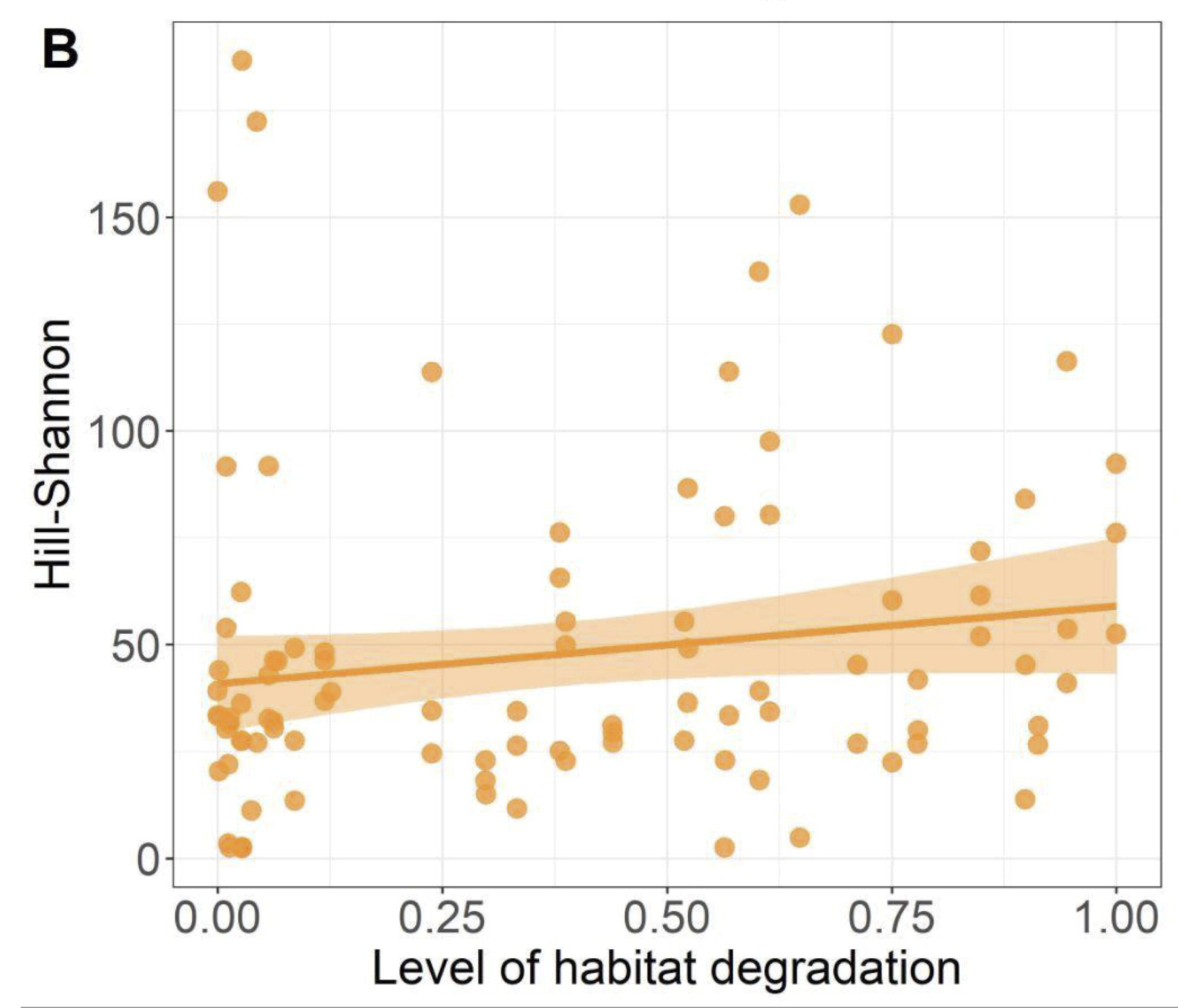
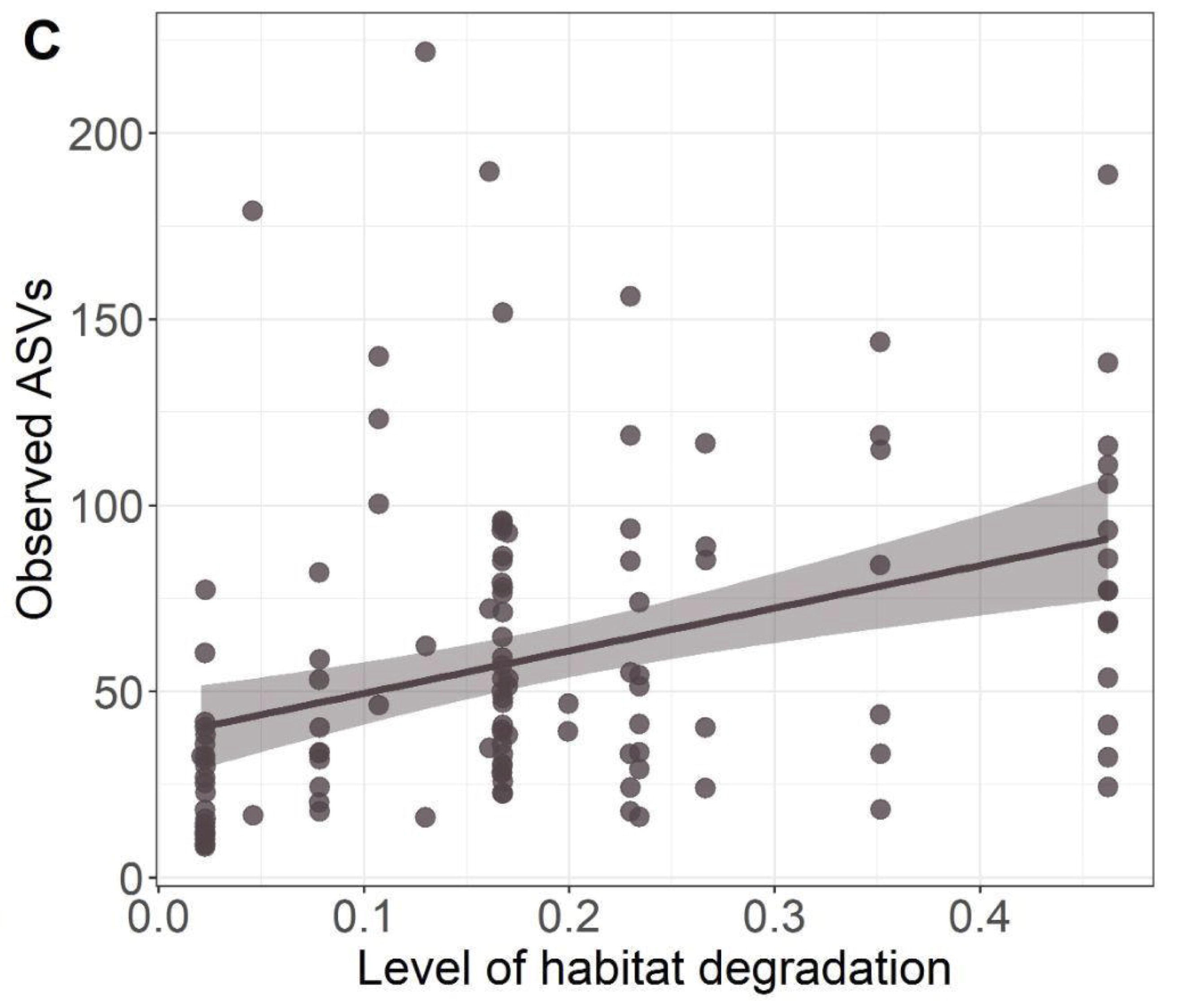
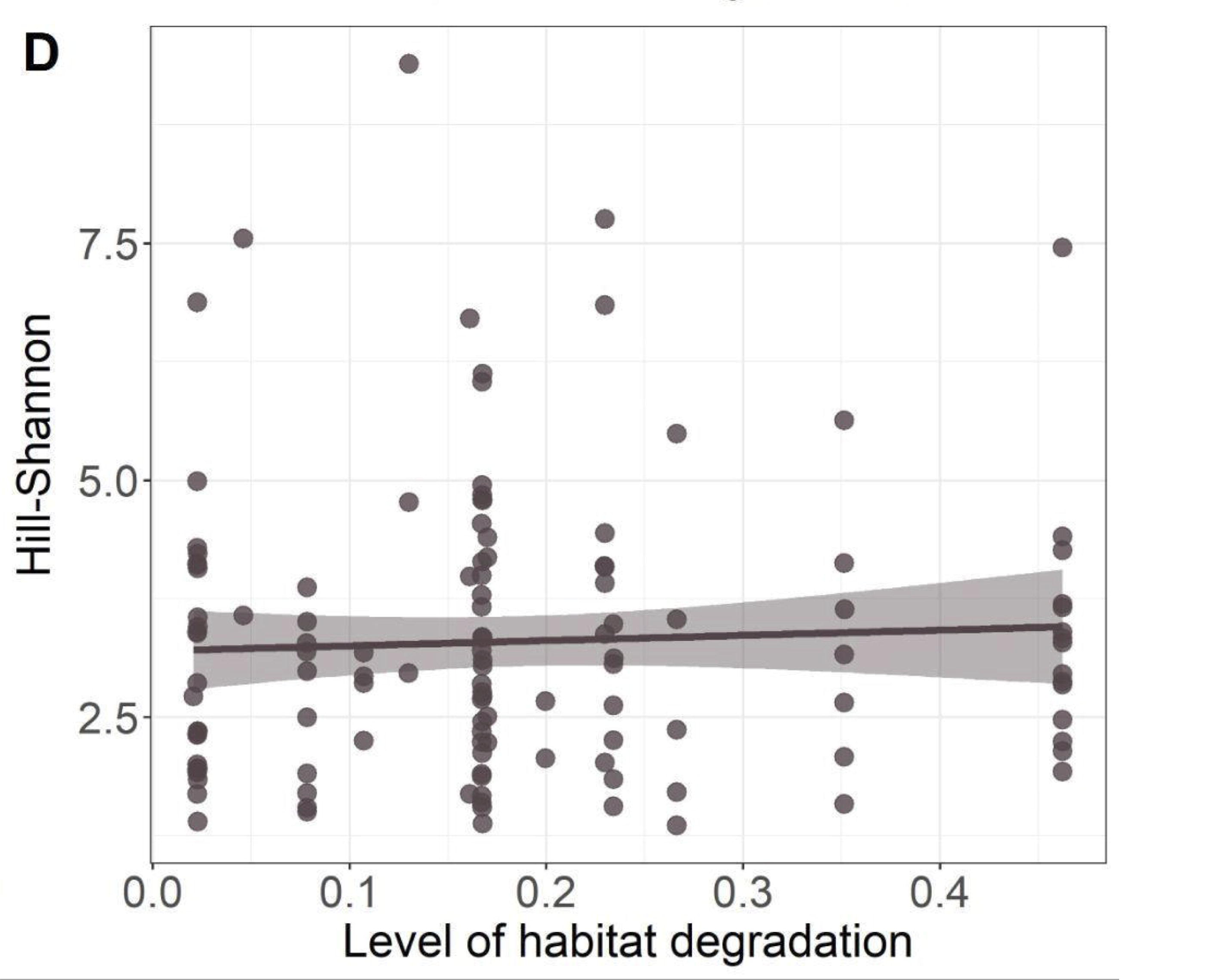
Variations in indices of alpha-diversity in the microbiota of the butterfly *M. cinxia* (orange) and the weevil *M. pascuorum* (grey) collected along a gradient of habitat degradation in a 10-meter buffer area surrounding each meadow. A) Observed ASVs and B) Hill-Shannon index for *M. cinxia*; and C) observed ASVs and D) Hill-Shannon index for *M. pascuorum*. Coloured areas along the lines indicate the 95% confidence intervals. *M. cinxia* were collected from a wider diversity of meadows across the Åland Islands than *M. pascuorum*.

For the weevil *M. pascuorum*, the level of degradation within habitat had no significant effect on the observed ASV count (*p*-value > 0.05; Fig. S3C). However, the level of degradation within habitat affected the Hill-Shannon estimates of alpha diversity (IRR = 11.92, 95% CI = 5.20 – 18.64, *p*-value = 0.001), with the Hill- Shannon index values significantly increasing with the level of degradation within habitat (Fig. S3D; Table S3). The level of habitat degradation in the 10m buffer surrounding each meadow had a significant effect on the observed ASVs (IRR = 11.87, 95% CI = 2.14 – 64.79, *p-*value = 0.01; Table S3), with the ASVs count increasing with the level of habitat degradation in the surrounding landscape (Fig. 4C). However, the level of habitat degradation in the 10m buffer surrounding each meadow did not significantly affect the Hill-Shannon estimates of alpha diversity (*p*-value > 0.05; Table S3; Fig. 4D). The focal meadow size had no effect on the observed ASV count, nor on the Hill-Shannon diversity of the microbiota (*p*-values > 0.05).

For *M. cinxia*, the indicator species analysis associated 12 bacterial ASVs to specimens collected from habitats surrounded by degraded habitats, and 20 ASVs to the more natural surroundings (Table S4). For *M. pascuorum*, the indicator species analysis suggested 114 ASVs to reflect the more degraded habitats, and 1 ASV belonging to the *Sodalis* genus to the more natural habitats (Table S4).

### Effects of habitat degradation on bacterial community composition (beta-diversity)

In *M. cinxia*, the level of habitat degradation within meadows had no effect on bacterial community composition (*p-*values>0.05; Fig. S4A &B). Habitat degradation in the 10m buffer surrounding the meadows did not influence the community composition Bray Curtis (Fig. 5A) but did influence the community composition Unweighted UniFrac (PERMANOVA: F = 2.156, R2 = 0.020, *p*-value = 0.009; Fig. 5B, Table S5), with a shift in community composition across the habitat degradation gradient. Sampling commune or focal meadow size had no effect on the composition of the microbiota of *M. cinxia* (*p*-value > 0.05). Most important bacterial ASVs affecting the PCoA biplot for Unweighted UniFrac were classified to the genus *Yersinia, Hymenobacter, Chrysobacterium, Sphigomonas* and phylum Proteobacteria; and for Bray Curtis, to genus *Yersinia, Wolbachia, Massilia, Chryseobacterium* and the phylum Proteobacteria (Fig. S5A & B).

**Figure 5.**
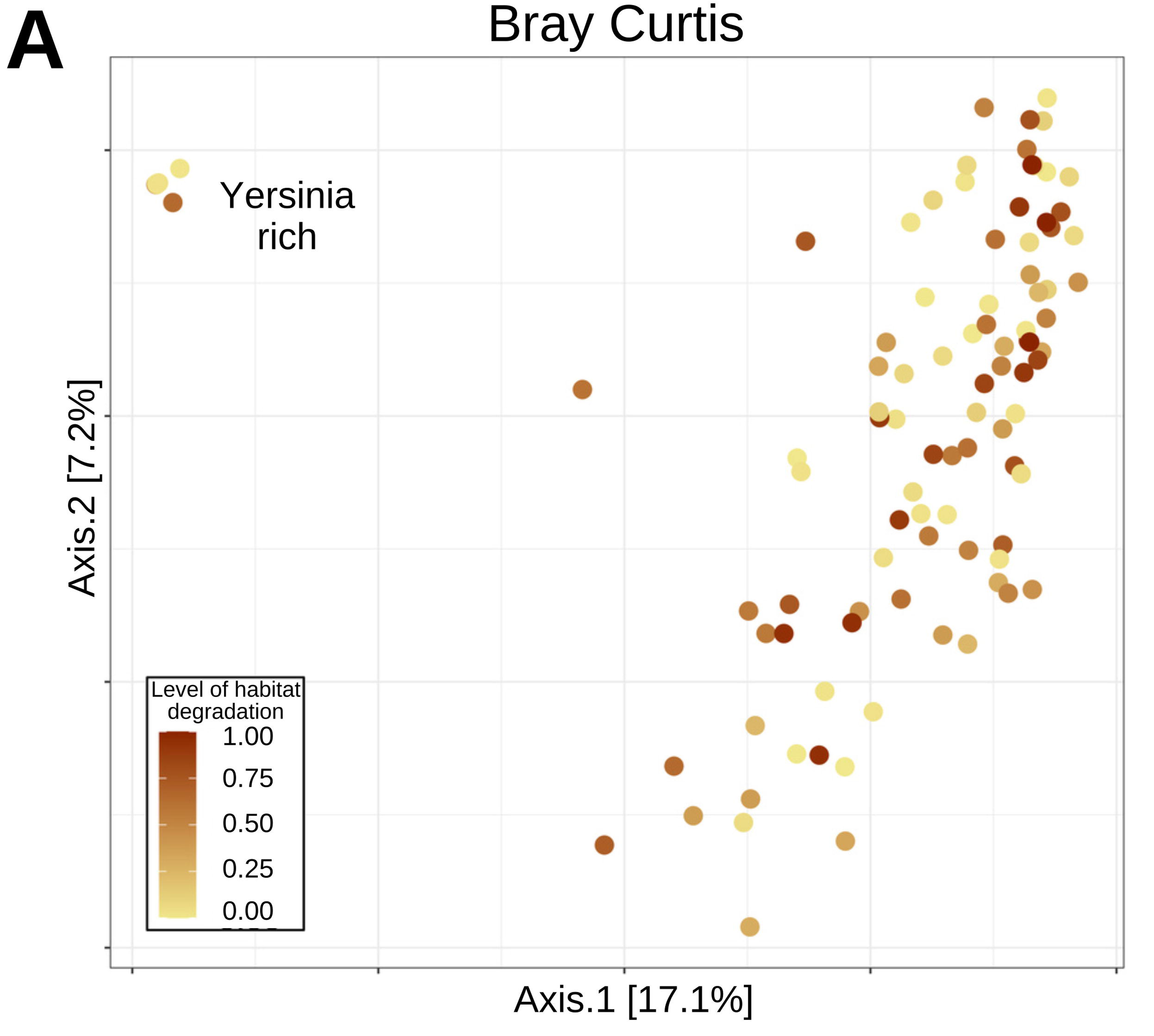
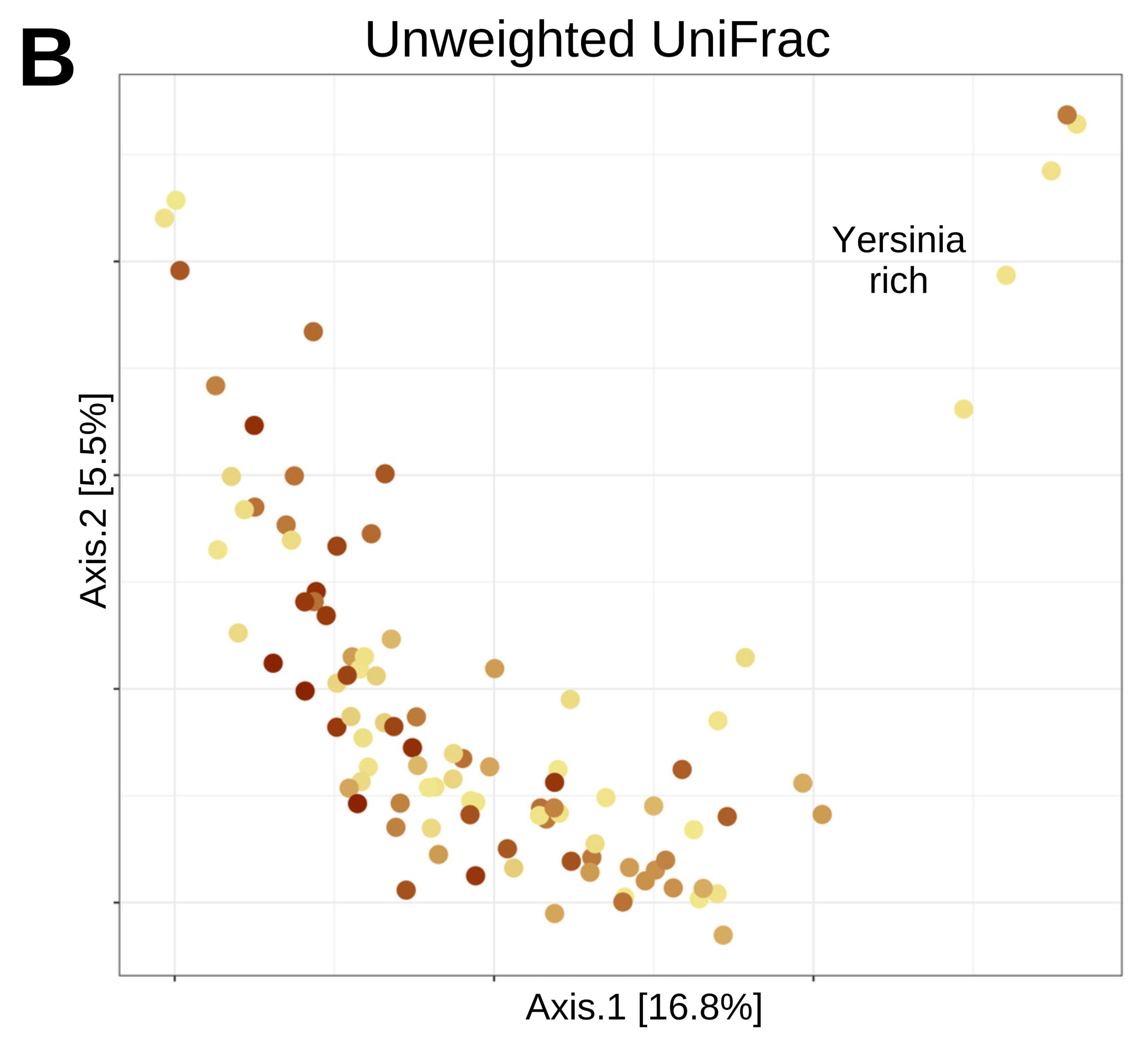
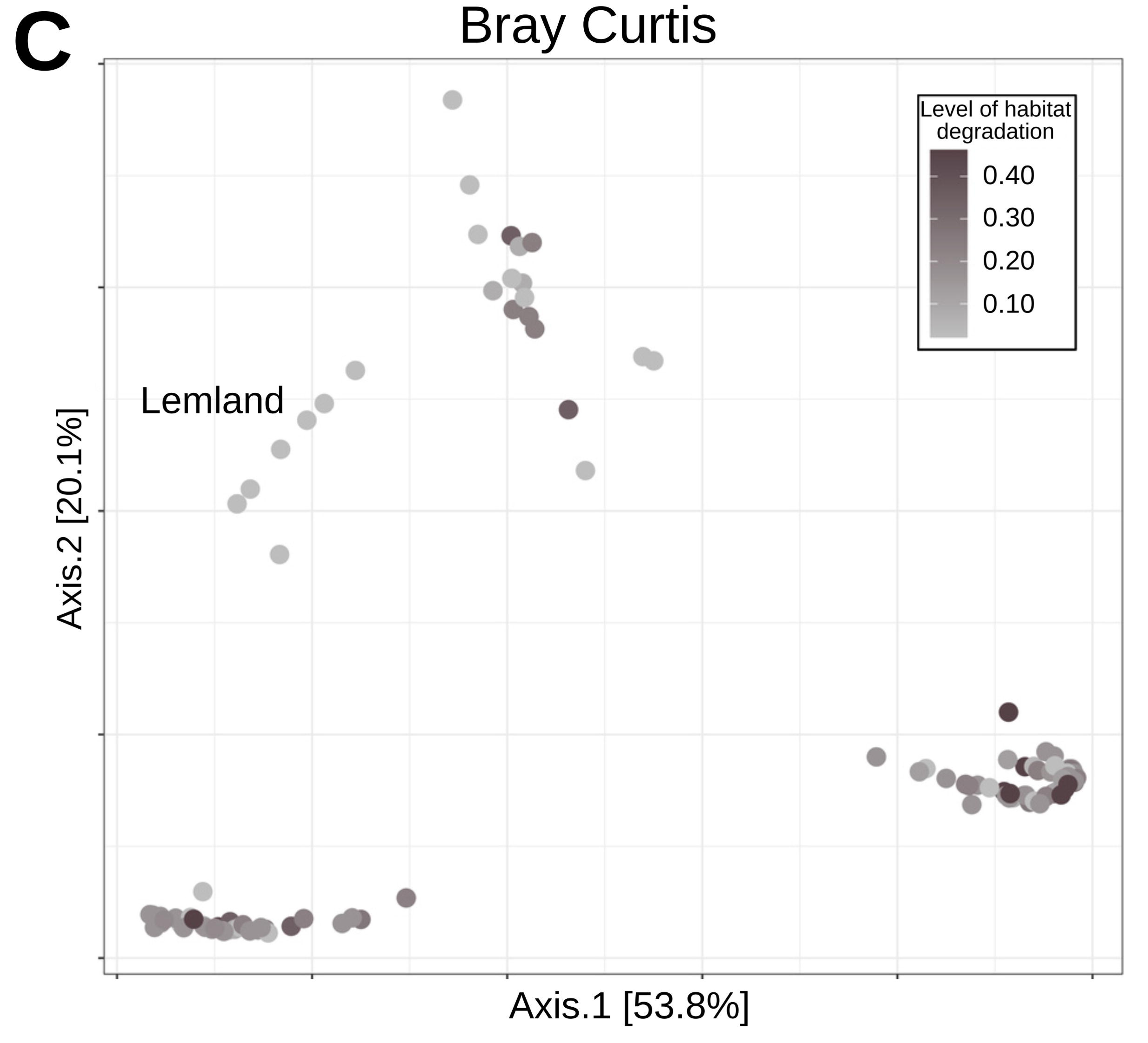
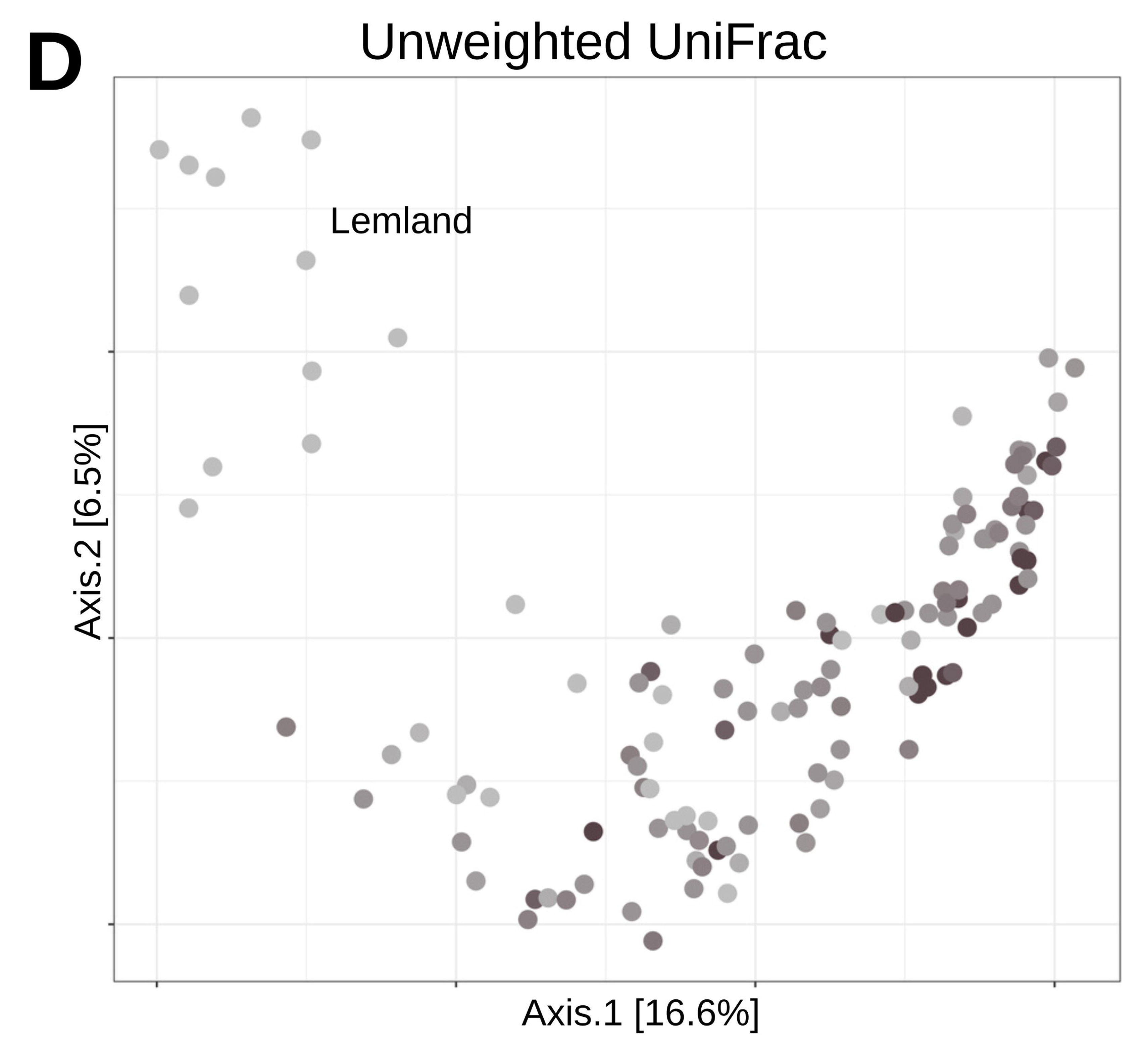
PCoA plots of the bacterial community composition (Beta-diversity) of the microbiota associated with (left) *M. cinxia* and (right) *M. pascuorum*, using either Bray Curtis dissimilarity (A & C) or Unweighted UniFrac (B & D) models. The colours darken as the level of habitat degradation in the 10-meter buffer area surrounding each meadow increases. Clear grouping of the ‘*Yersinia-rich*’ *M. cinxia* specimens, and ‘*Lemland*’ weevil specimens, are indicated in each panel.

In *M. pascuorum*, the bacterial community composition was affected by the level of habitat degradation. Both Bray Curtis and Unweighted UniFrac PERMANOVA models suggest that the composition of the microbiota was impacted by both the level of degradation within meadow (Bray Curtis: PERMANOVA: F = 5.37; R2 = 0.034; *p*-value = 0.005; Unweighted UniFrac: PERMANOVA: F = 2.062; R2 = 0.015; *p*-value = 0.007; Fig. S4 C & D; Table S5), and by the level of habitat degradation in the 10m buffer surrounding the meadows (Unweighted UniFrac: PERMANOVA, F = 6.760, R2 = 0.049, *p*-value = 0.001, Fig. 5C; Bray Curtis: PERMANOVA, F = 10.499, R2 = 0.066, *p*-value = 0.001, Fig 5D; Table S5). The specimens from more degraded habitats carry more homogenous microbiota. The PCoA plot clearly suggests that the microbiota of *M. pascuorum* weevils form four different groups, and sampling commune affected the community composition of the microbiota of *M. pascuorum* (Bray Curtis: PERMANOVA, F = 3,459; R2 = 0.154; *p*-value = 0.001; Unweighted Unifrac: PERMANOVA, F = 1.761; R2 = 0.090; *p*-value = 0.001) (Table S5) with specimens from Lemland showing a community profile enriched with *Sodalis sp*. (Fig. S6A & B), a mutualistic bacterium providing threonine to weevils (Vigneron et al 2014). Most important ASVs affecting the PCoA biplots were classified to the genera *Sodalis* and *Rickettsia*, and to the family Morganellacaea (Fig. S6A & B). Finally, the size of the meadow influences the community composition of the microbiota associated with the weevils (Bray Curtis: PERMANOVA, F = 3.480, R2 = 0.022, *p*-value = 0.019; Unweighted Unifrac: PERMANOVA, F = 2.487, R2 = 0.018, *p*-value = 0.002).

### Effects of habitat degradation on the predicted functionality of insect-associated bacterial communities

The predicted functionality analyses revealed no effect of the level of habitat degradation (*p*-values > 0.05, Fig. 6A, Table S6), and no effect of the focal meadow size (*p*–value > 0.05) on the microbial predicted functionality in the butterfly *M. cinxia*. Sampling commune affected the predicted functionality of the microbiota in *M. cinxia* (PERMANOVA: F = 1.382, R2 = 0.082, *p*-value = 0.006). In contrast, there was an effect of the level of habitat degradation (within meadow: PERMANOVA: F = 30.391, R2 = 0.174, *p*-value = 0.001; 10m buffer surrounding the meadow: PERMANOVA: F = 9.197; R2 = 0.053, *p*-value = 0.002), sampling commune (F = 2.235; R2 = 0.095, *p*-value = 0.012) with the microbiota of weevils from the Lemland commune being functionally different because rich in *Sodalis* symbionts. However, there was no effect of focal meadow size (*p*-value > 0.05) on the predicted functionality of the microbiota of the weevil *M. pascuorum* (Fig. 6B, Table S6).

**Figure 6.**
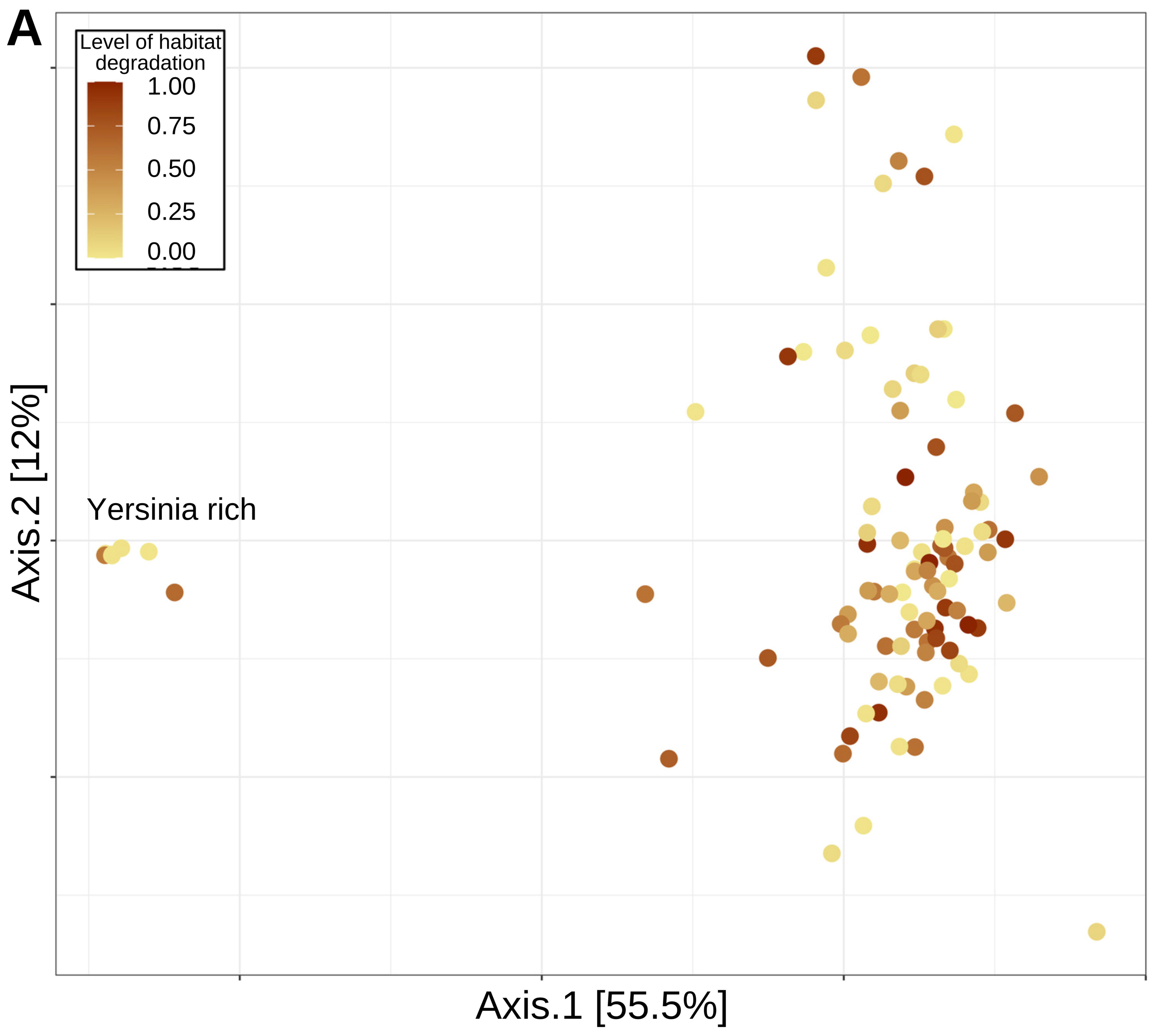
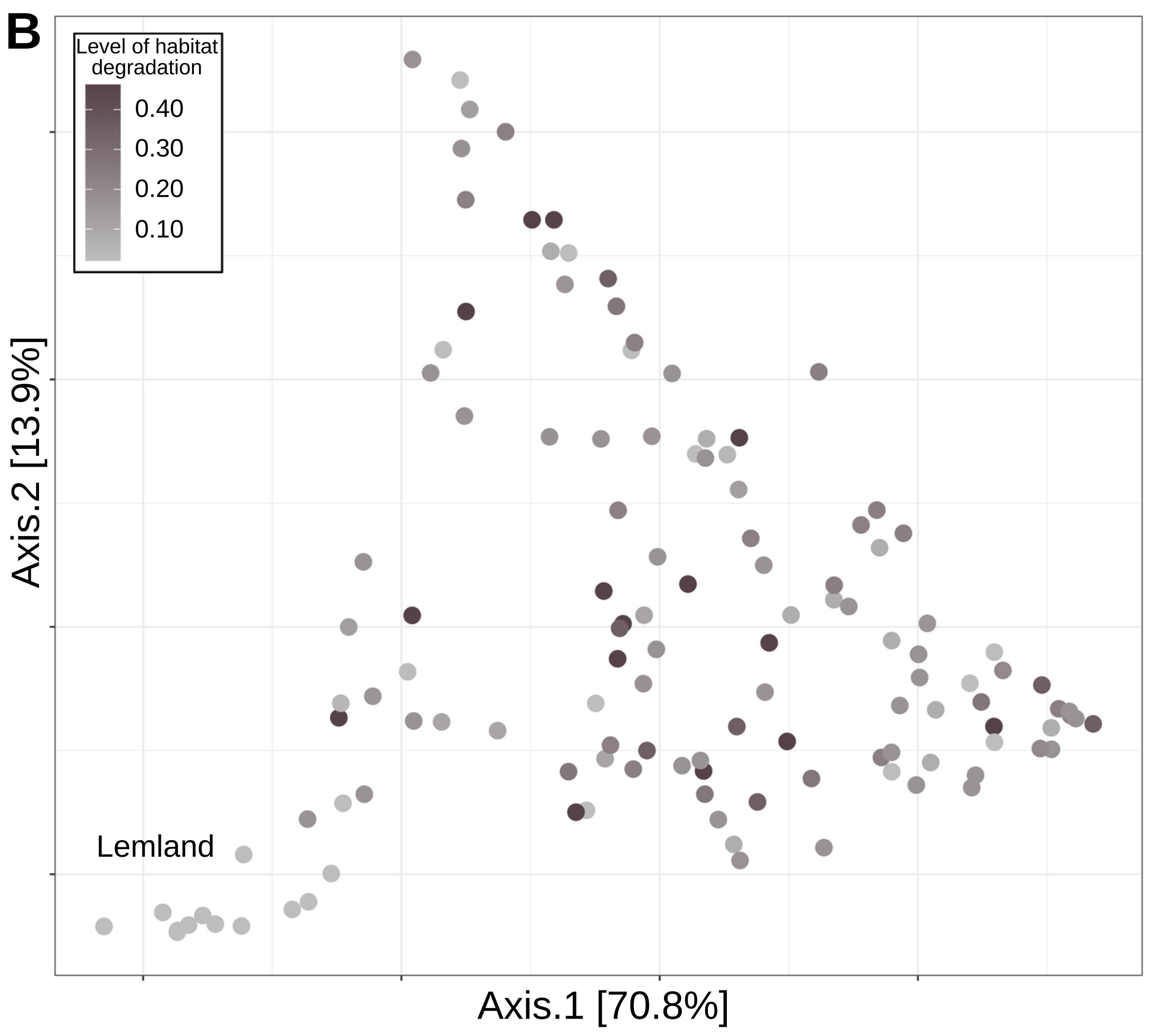
PCoA plots for the q2-picrust2 produced predicted metabolic pathway analyses using Bray Curtis distances for A) *M. cinxia* and B) *M. pascuorum.* Visually clear grouping of the ‘*Yersinia-rich*’ caterpillars, and weevils from the ‘*Lemland commune*’ are indicated in either relevant figure panel.

## Discussion

Although both the caterpillars of the butterfly *M. cinxia* and the *M. pascuorum* adult weevils feed on the leaves of the same host plant species, *P. lanceolata* (Nieminen and Vikberg 2015, Ojanen et al 2013), here we show that their respective associated bacterial communities are very different. *Melitaea cinxia* carry a complex microbiota, rich in environmental bacterial species, which have no to little effects on their butterfly host biology (Duplouy et al 2020, Minard et al 2019). Instead, the microbiota of the weevil quite strikingly convened into four distinct groups, each dominated by a different maternally inherited symbiotic bacterium (ie. two *Rickettsia, Sodalis* and *Wolbachia*), and our microbial community models could only predict the microbiota profile of the weevil host. These results support the idea that the microbiota of *M. cinxia* butterflies is colonized by a transient microbiota subject to stochastic processes, while that of the weevils is more predictable because resident and mutualistic, and thus subject to deterministic processes (Anbutsu et al 2017, Jing et al 2020, Masson et al 2015, Muhammad et al 2019, White et al 2015).

Resident microbiota are often dominated by inherited bacteria, which have been shown to manipulate their host reproductive system (Giorgini et al 2010, Hsiao and Hsiao 1985, Perlman et al 2006), support egg production (Perotti et al 2006), contribute to their host resistance against parasites (White et al 2015, Łukasik et al 2013) or response to heat stress (Tougeron and Iltis 2022). The weevils from the Lemland commune in the Åland Islands group together as their microbiota is abundant with *Sodalis* bacteria. Vigneron et al. (2014) showed that in the cereal weevil *Sitophilus*, *Sodalis* densities increases within a week of the weevils moulting but decreases once the new cuticle has hardened. *Sodalis* are maternally inherited symbiotic bacteria, which provide weevils with amino-acid necessary to the production of their thick protective cuticle after molting (Anbutsu et al 2017). Unfortunately, as our study does not include any trait and fitness measures for the host species, it remains unclear whether the dispersal ability, health or any other fitness traits of the host may vary with their associated microbial communities. Future studies following changes in the density of these symbionts will inform on the exact nature of the symbioses under different environmental and stress conditions.

In this context, our study of the microbiota of both insects provide evidence that the degradation of the host’s habitat under anthropogenic activities can modify local species community (Decaëns et al 2018, Strona and Lafferty 2016) even at the microbial level. Even though the level of habitat degradation within the meadow, and the size of the focal meadow, only affect the bacterial community composition associated with the weevil *M. pascuorum*, in both species, the level of landscape degradation (ie. land-use changes in the 10m buffer surrounding the meadows) leads to a shift in the species composition of their associated bacterial microbiota. This results occurs in both species, even though they were collected on different years, and from a different set of meadows, suggesting a potentially conserved effect of habitat degradation across species and time in this system. However, in contrast with our original expectations, habitat degradation through land-use changes and reduction of habitat size, does not lead, in our system, to the loss of functional bacterial species diversity. Rather the opposite, we show that with habitat degradation the bacterial species richness indices remain unchanged in the butterfly *M. cinxia* and even significantly increase in the weevil *M. pascuorum*. As microbial diversity losses are often more characteristic of stressed specimens (Wei et al 2017), this results might suggest that despite habitat degradation, the Åland Islands are inhabited by an insect community with a healthy microbiota resistant to disturbances (Lozupone et al 2012).

For many species, the composition of the microbiota, and especially that of the gut microbiota, is representative of their diet (McManus et al 2021). Recently, Teng et al (2022) suggested that differences in the gut microbiota associated with populations from three rodent species were linked to changes in diets between populations. The animals from farmlands, where food diversity is high, hosted the most species- rich microbiota (Teng et al 2022). In Åland, both the adult weevils and the caterpillars likely remain within their own meadow (Smith et al 2021), where the weevil *M. pascuorum* is a specialist herbivore of *P. lanceolata*, feeding on the leaves as adults, and on the seeds as larvae (Nieminen and Vikberg 2015), while, the caterpillars of *M. cinxia* can feed on a second host plant *Veronica spicata*, which was not recorded within our selected meadows (Hanski et al 2002, Hanski 2011). The increased bacterial diversity in *M. pascuorum* specimens collected from more degraded habitats is thus unlikely due to these insects switching diet or feeding on a wider diversity of host plants in the meadows degraded by human activities. Alternatively, the habitat degradation could lead to changes in the host plant, the host plant associated microbiota, or in other element of the insects’ environments. *Plantago-*associated bacterial communities are indeed known to vary with the road network in the Åland Islands (Numminen and Laine 2020), but also with the abundance of defensive iridoid glycoside compounds within the host plant (Minard et al 2022), and with local soil microbial composition (Mursinoff and Tack 2017). Changes to local soil microbiota due to habitat degradation have been characterized worldwide (Kiesewetter and Afkhami 2021, Li et al 2019), and as the weevils crawl on soil surfaces when moving between plants and plant leaves (Hannula et al 2019), they could collect microbial species from their local soil.

Another explanation for the differences we found in microbial communities among taxa could be that habitat degradation allows for the spread of diseases, which can disturb the stability of microbial communities, and allow the colonization of immune-challenged specimens by new microbes (Abraham et al 2017, Fredensborg et al 2020, Zhang et al 2016). The detection of *Yersinia* bacteria in few *M. cinxia* caterpillars may suggest that our sampling included sick individuals upon collection in the field as some *Yersinia* species are toxic to animals including invertebrates (Bresolin et al 2006, Springer et al 2018). In *M. cinxia* caterpillars, the clear shift towards a microbiota dominated by Enterobacteriaceae bacteria was previously observed in field collected *M. cinxia* specimens (Minard et al 2019), but not in caterpillars reared in the laboratory (Duplouy et al 2018, Duplouy et al 2020). In contrast, the abundance of the bacterial taxon *Pseudomonas* in the transient gut microbiota of caterpillars from natural habitats may simply indicate that the bacterium is more abundant in these habitats, the cause of which remains unknown. *Pseudomonas* bacteria colonize a wide range of niches, some species are ubiquitous from soil, plant and the freshwater environments, some are opportunistic pathogens of humans and other organisms (Moore et al 2006), including insects (Flury et al 2016).

Although habitat degradation is often associated with negative effects on macro-biodiversity level (Soh et al 2019, Teng et al 2022, Uhl et al 2021), the study of the impact of such Anthropogenic changes on microbial communities associated with species (ie. symbioses) remains scarce (Hom and Penn 2021). Nonetheless, evaluating the responses of symbiotic microbial communities to the habitat changes is crucial to predicting the vulnerability and resilience of macro-species to habitat degradation and any other anthropogenic changes. Our study contributes to bridging this gap by showing that degradation of the host habitat through land-use changes can indeed induce a shift in the composition of microbial communities, and even have a positive effect, on the overall bacterial species richness associated with insects. This is in clear contradiction with our original expectations that degradation of the host habitat would also be detrimental to their associated microbial diversity and functionality (McManus et al 2021, Teng et al 2022). One possible explanation for these results is that habitat degradation in the Åland meadow network is not strong enough to lead to microbial erosion in the insect species studied. Rather it seems to offer opportunities for the insects to potentially experience more heterogeneous habitats, which could support different or more diverse sets of microbial species with similar functionality for their host. Although our system does not allow to test stronger levels of habitat degradation, it still suggests that remaining at least below a threshold of habitat degradation could support the functional stability of the microbiota associated with insects. The exact character of that habitat degradation threshold however remains to be further identified.

## Data accessibility

The metabarcoding raw sequence are available from the NCBI database under the NCBI SRA project number PRJNA912103.

## Supporting information

Sup materiel Tables

## Acknowledgements

We would like to thank Prof. S. van Nouhuys, Dr. M. Nieminen, all the members of the ISEE and life-history evolution groups for discussions on the study. Thanks to Dr. T. Schulz for guidance with the land-use data, and to T. Hannunen from the Finnish Institute for Molecular Medicine for the sequencing of the *16S* srRNA amplicons. We wish to acknowledge CSC – IT Center for Science, Finland, for computational resources. Also, thanks to the yearly Åland survey participants for collecting the caterpillars in the field, and to the Lammi Biological Station for providing facilities to keep diapausing caterpillars.

