## Supplementary material for "Species-associated bacterial diversity increases along a gradient of habitat degradation": Sup material Figures

**Supplementary figures**

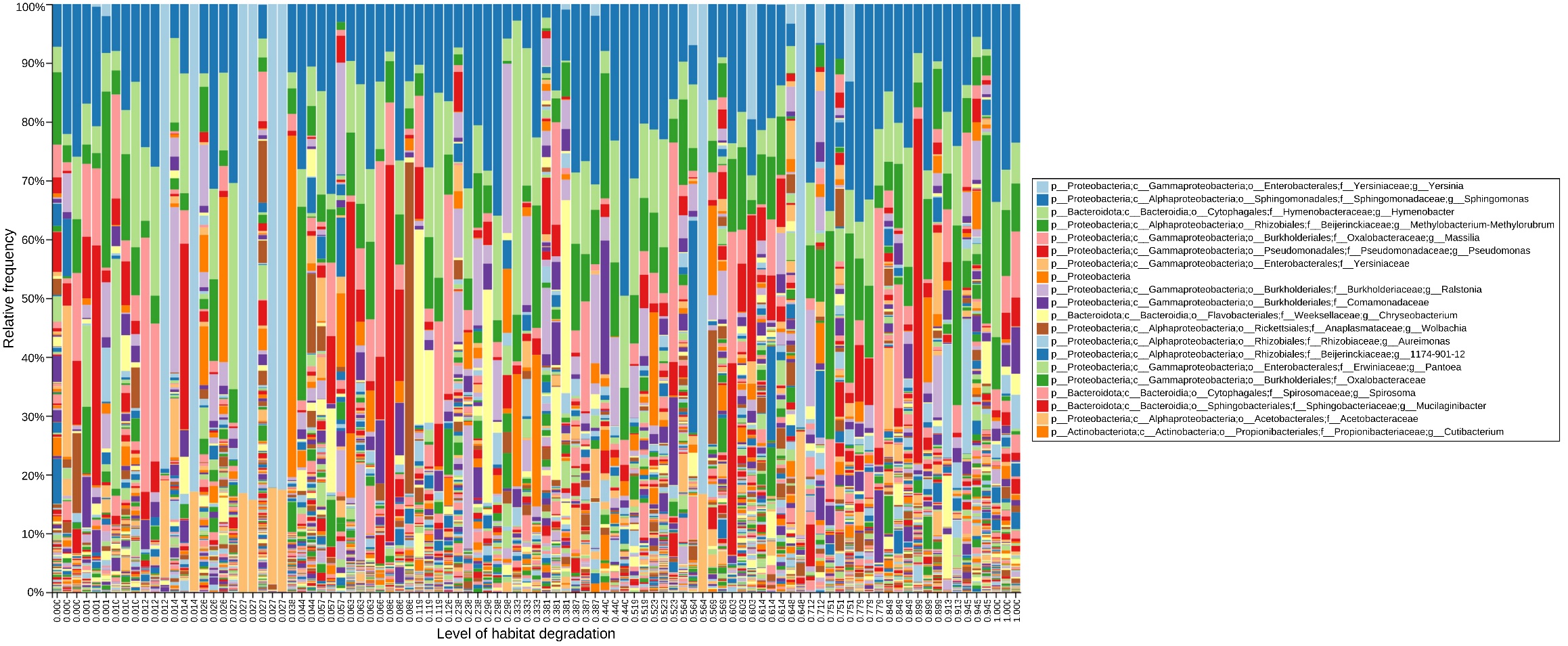

**Supplementary Figure 1A**. **Relative frequencies of each genus in the *M. cinxia* microbiota samples**, with top 20 genera visible in the legend. Each bar represents a separate sample, with sample labels indicating the proportion of habitat degradation in the 10-meter buffer surrounding each sampling patch.

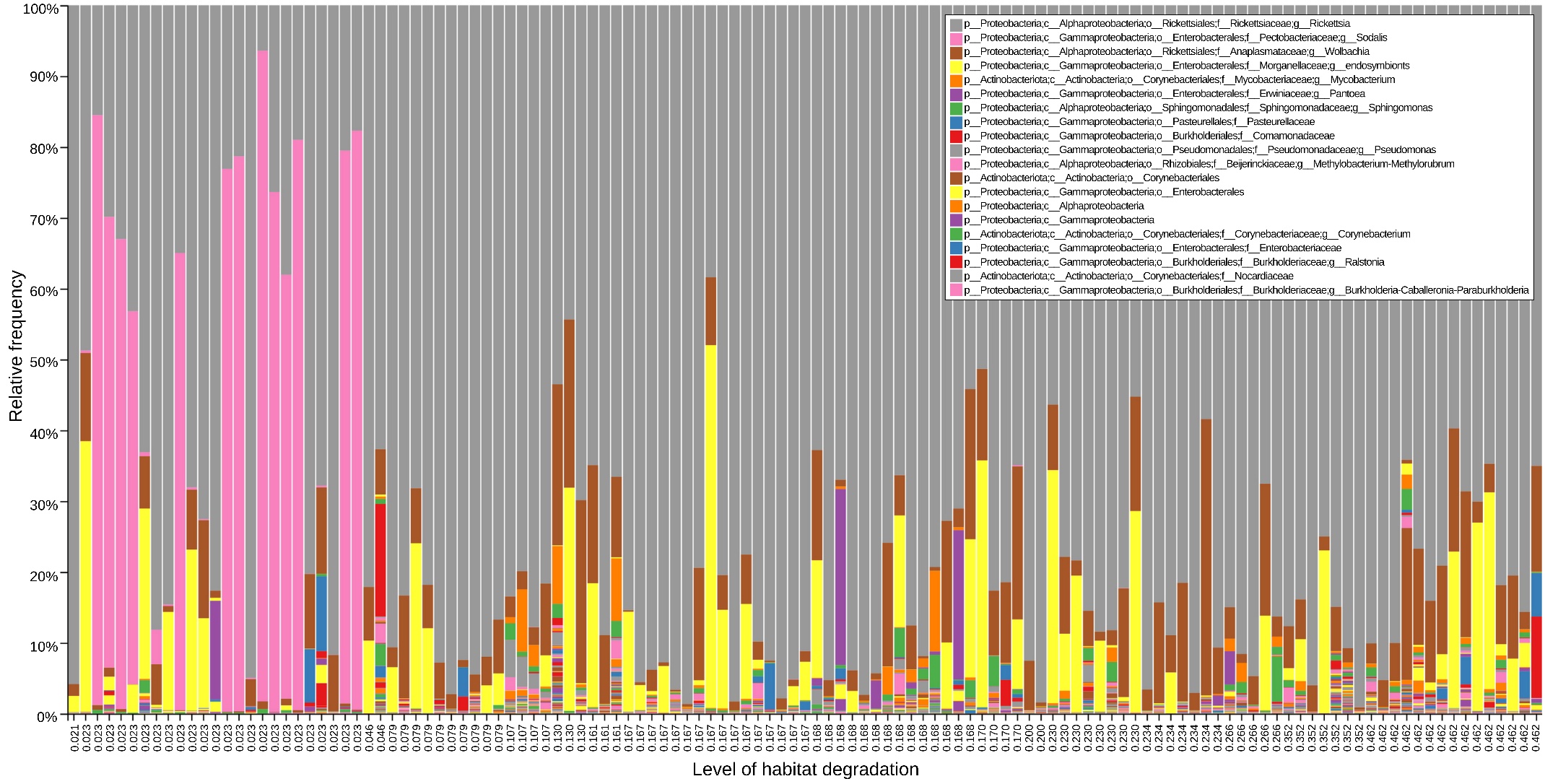

**Supplementary Figure 2B. Relative frequencies of each genus in the *M. pascuorum* microbiota samples**, with top 20 genera visible in the legend. Each bar represents a separate sample, with sample labels indicating the proportion of habitat degradation in the 10-meter buffer surrounding each sampling patch.

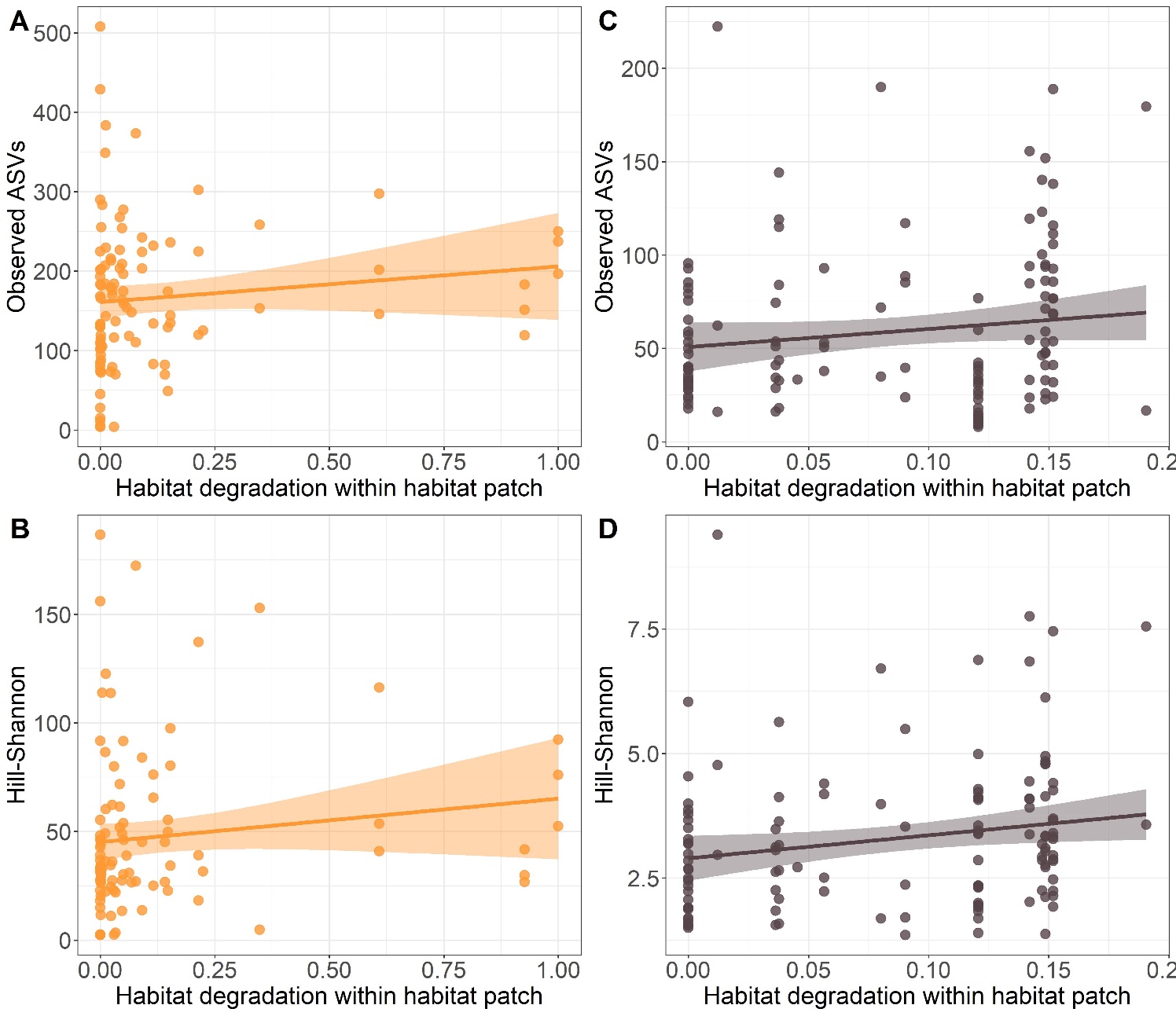

**Supplementary Figure 3. Variation in bacterial alpha-diversity, as A&C) observed ASV’ and B&D) Hill-Shannon index associated with either *M. cinxia* (orange) and *M. pascuorum* (grey) along a gradient of habitat degradation within patches (rather than in the surrounding habitats).** Coloured areas along the lines indicate the 95% confidence intervals. *M. cinxia* and *M. pascuorum* were collected from different habitat patches across the Åland island. Habitats sampled for the *M. cinxia* had more variability in the level of degradation.

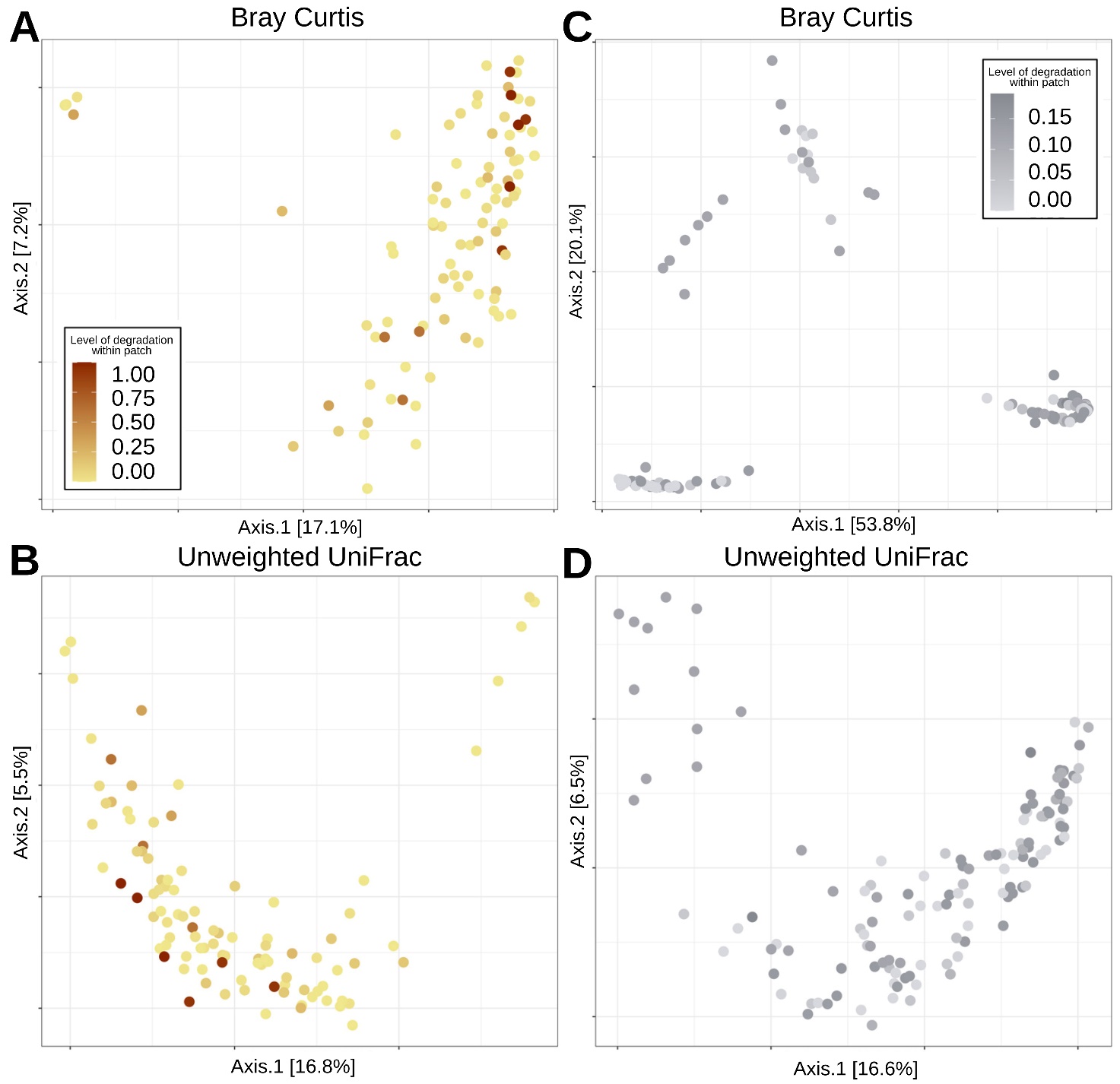

**Supplementary Figure 4. PCoA plots for the composition of bacterial communities (Beta-diversity) for (left) *M. cinxia*’s and (right) *M. pascuorum*’s associated microbiota**, using either Bray Curtis dissimilarity (A & C) or Unweighted UniFrac (B & D) methods. The colours darken as the proportion of habitat degradation within patches increases.

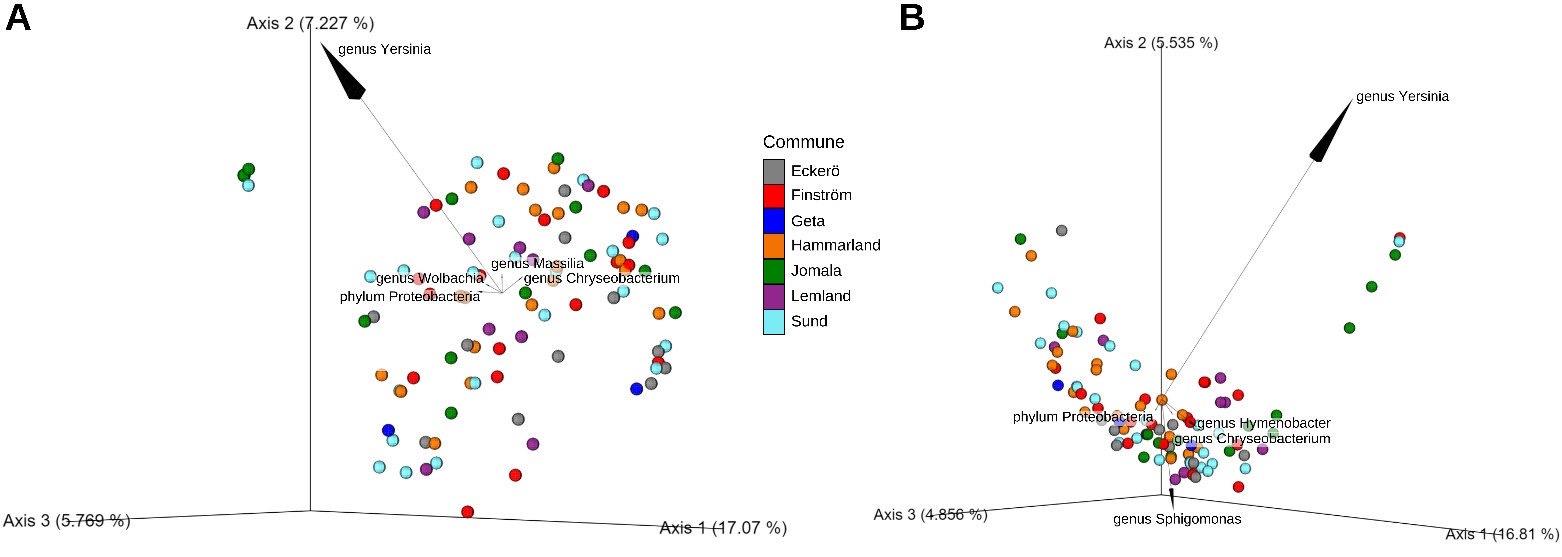

**Supplementary Figure 5.** **Biplots indicating the 5 most important bacterial ASVs affecting the PCoA plots of the butterfly *M. cinxia* microbiota for a) Bray Curtis dissimilarity and b) unweighted UniFrac.** The bacterial genus (or if not available, the most precise classification available) is indicated on the figure next to a corresponding arrow. Individual samples are coloured according to the commune where the sample was collected from. The plots are presented as 3D plots of the first three PCoAs.

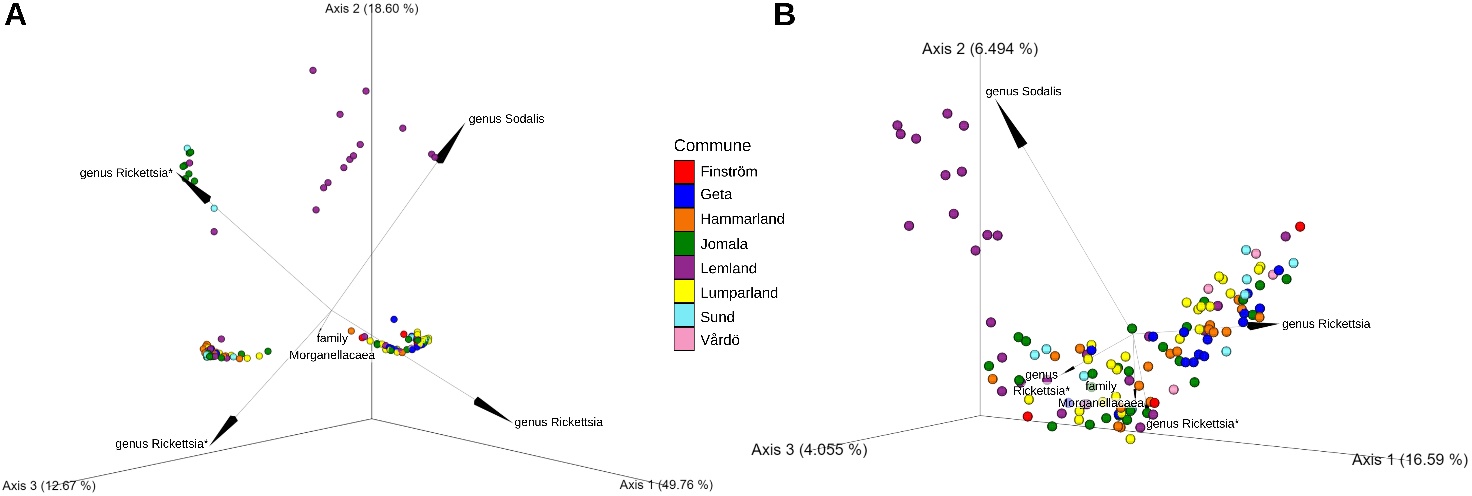

**Supplementary Figure 6. Biplots indicating the 5 most important bacterial ASVs affecting the PCoA plots of the weevil *M. pascuorum* microbiota for a) Bray Curtis dissimilarity and b) unweighted UniFrac**. The bacterial genus (or if not available, the most precise classification available) is indicated on the figure next to a corresponding arrow. Individual samples are coloured according to the commune where the sample was collected from. The symbol (*) next to two Rickettsia indicates classification to the same species (*Rickettsia bellii*). The plots are presented as 3D plots of the first three PCoAs.
